# mRNA condensation fluidizes the cytoplasm

**DOI:** 10.1101/2023.05.30.542963

**Authors:** Ying Xie, Tiewei Liu, David Gresham, Liam J Holt

## Abstract

The intracellular environment is packed with macromolecules of mesoscale size, and this crowded milieu significantly influences cell physiology. When exposed to stress, mRNAs released after translational arrest condense with RNA binding proteins, resulting in the formation of membraneless RNA protein (RNP) condensates known as processing bodies (P-bodies) and stress granules (SGs). However, the impact of the assembly of these condensates on the biophysical properties of the crowded cytoplasmic environment remains unclear. Here, we find that upon exposure to stress, polysome collapse and condensation of mRNAs increases mesoscale particle diffusivity in the cytoplasm. Increased mesoscale diffusivity is required for the efficient formation of Q-bodies, membraneless organelles that coordinate degradation of misfolded peptides that accumulate during stress. Additionally, we demonstrate that polysome collapse and stress granule formation has a similar effect in mammalian cells, fluidizing the cytoplasm at the mesoscale. We find that synthetic, light-induced RNA condensation is sufficient to fluidize the cytoplasm, demonstrating a causal effect of RNA condensation. Together, our work reveals a new functional role for stress-induced translation inhibition and formation of RNP condensates in modulating the physical properties of the cytoplasm to effectively respond to stressful conditions.

## Introduction

The cytoplasm undergoes rapid changes in its physical properties in response to stress. Notably, when exposed to ATP depletion by chemicals or glucose starvation, the cytoplasm of bacterial and yeast cells can undergo a solid-state transition characterized by a significant reduction in macromolecular motion (Joyner et al., 2016; Munder et al., 2016; Parry et al., 2014). However, an immediate and uniform slowdown of all cellular processes might not be helpful in all stress conditions, since there are many processes that must be coordinated in a time-dependent manner to achieve stress adaptation. For example, stress responsive transcription factors translocate from the cytosol to the nucleus within minutes of *Saccharomyces cerevisiae* being exposed to environmental stresses, activating stress response transcription (Delaunay et al., 2000; Görner et al., 2002; Muzzey et al., 2009; Triandafillou et al., 2020). Also, the cell significantly rearranges plasma membrane (Laidlaw et al., 2021), vacuole membrane (Seo et al., 2017) and interorganelle membranes such as nuclear-vacuole junction (Wood et al., 2020) in response to glucose starvation. Finally, new structures form within the cytoplasm upon stress. Some misfolded proteins are directed to Q-bodies, which are membraneless organelles that form upon heat stress and glucose starvation (Escusa-Toret et al., 2013; Kaganovich et al., 2008; Sathyanarayanan et al., 2020; Sontag et al., 2022). Formation of Q-bodies depends on the interaction between the HSP42 chaperone and misfolded proteins (Specht et al., 2011). The large-scale remodeling of multiple cellular processes required in response to stress is incompatible with an immediate solidification of the cytoplasm. Therefore, more detailed studies of the physical properties of the cytoplasm during the initial response to stress are necessary.

Cells are highly crowded with mesoscale (20 nm-1 µm diameter) macromolecules (Ekman et al., 2017) (Figure 1A). Active biological processes can influence the motion of mesoscale macromolecules. For example, the dynamic assembly of microtubules enhances mesoscale diffusion within the densely packed metaphase spindle (Carlini et al., 2020). This crowded, active environment is crucial for efficient cellular functions, but the mechanisms controlling these biophysical factors and their influence on biochemistry are only beginning to be understood.

**Figure 1.**
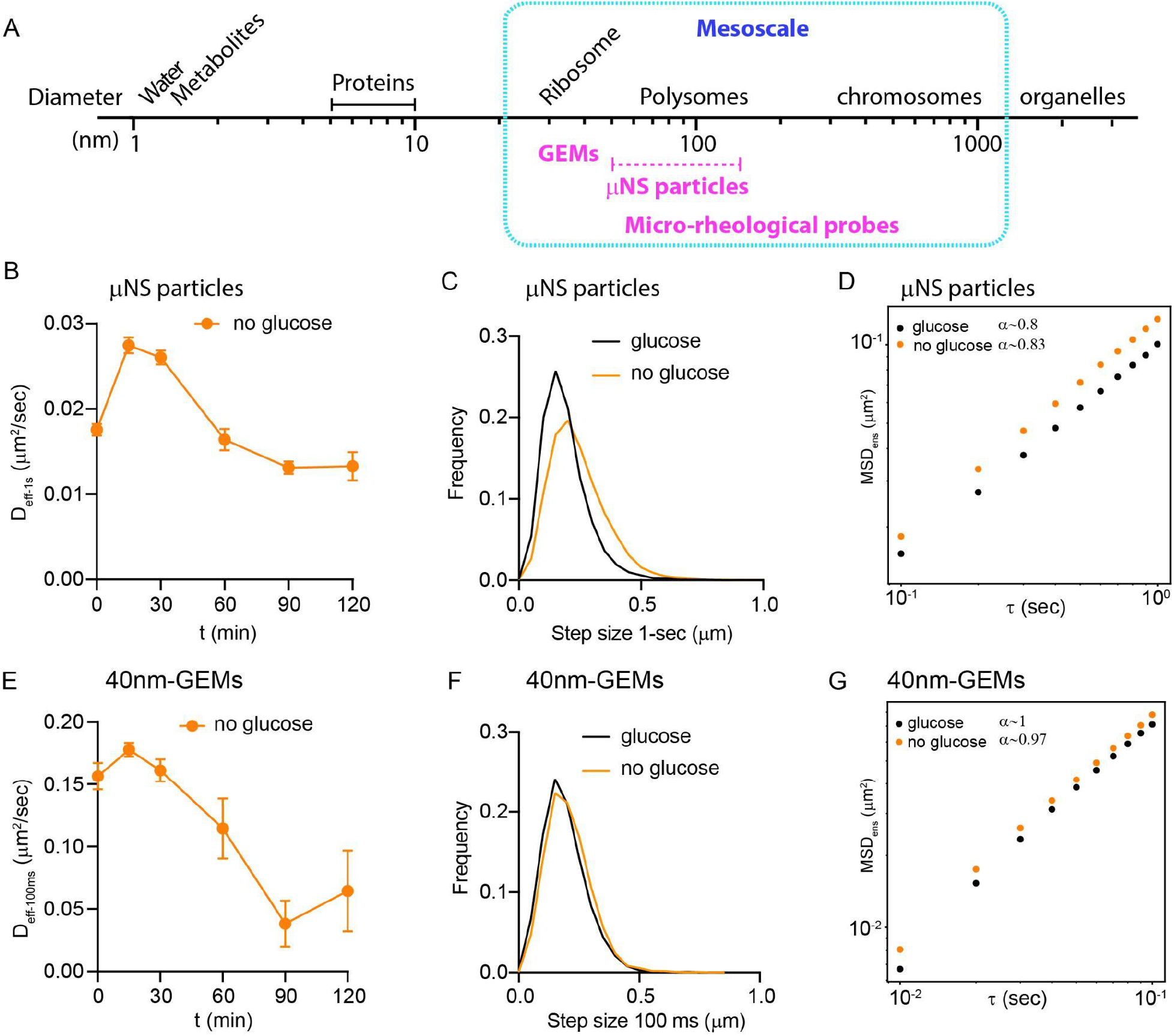
Acute glucose starvation leads to a transient increase in diffusivity of mesoscale macromolecules. **A.** Mesoscale rheological probe sizes in relation to macromolecular complexes. **B.** The effective diffusion coefficients at one second (D_eff-1s_) for μNS particles from 3 biological replicate experiments upon acute glucose starvation in *S. cerevisiae* cells (Median values are used to summarize each biological replicate. The mean and SEM of 3 biological replicates are shown). **C.** Step size distribution of μNS particles in one second time scale in glucose rich (glucose) or glucose depleted (no glucose) media at 15 minutes (glucose: n=10,437 trajectories; no glucose: n=16,372 trajectories). **D.** Ensemble-averaged mean-squared displacement (MSD) versus time delay (τ), log_10_ scale. A linear model was fit to determine the anomalous exponent α values for glucose rich (glucose) or glucose depleted (no glucose) condition at 15 minutes (glucose: n=10,437 trajectories; no glucose: n=16,372 trajectories). **E.** The effective diffusion coefficients at 100 msec (D_eff-100ms_) for 40nm-GEMs from 3 biological replicate experiments upon acute glucose starvation in *S. cerevisiae* cells (Median values are used to summarize each biological replicate. The mean and SEM of 3 biological replicates are shown.). **F.** Step size distribution of 40nm-GEMs in 100 msec time scale in glucose rich (glucose) or glucose depleted (no glucose) media at 15 min (glucose: n=6382 trajectories; no glucose: n=5116 trajectories). **G.** Ensemble-averaged mean-squared displacement (MSD) versus time delay (τ), log10 scale. A linear model was fitted to determine the anomalous exponent α values for glucose rich (glucose) or glucose depleted (no glucose) condition at 15 min (glucose: n=6382 trajectories; no glucose: n=5116 trajectories)

In a previous study, ribosomes, with a diameter of 25 nm, were identified as dominant mesoscale macromolecular crowders in the cytoplasm, accounting for approximately half of the total excluded volume (around 20% of the total cytoplasmic volume) (Delarue et al., 2018). Inhibition of the TORC1 kinase reduced the concentration of ribosomes, thereby decrowding the cytoplasm, which both increased the diffusivity of mesoscale macromolecules and influenced the formation of phase-separated condensates (Delarue et al., 2018). Ribosomes are further organized into translating polysome complexes, which vary in size depending on the number of ribosomes bound to the mRNA. These larger cytoplasmic structures actively undergo rearrangement and can create “local cages” that restrict mesoscale diffusivity, confining larger macromolecules while allowing smaller molecules to diffuse more easily through these local cages (Parry et al., 2014; Xiang et al., 2021).

In response to various environmental stresses, there is a general suppression of protein production, primarily by inhibiting the rate-limiting step of translation initiation (Advani and Ivanov, 2019; Crawford and Pavitt, 2019; Janapala et al., 2019). In yeast, glucose starvation triggers a significant reduction of translation within minutes, leading to disassembly of polysome complexes into monosomes, and the release of mRNA (Ashe et al., 2000; Bresson et al., 2020; Kuhn et al., 2001). The mRNA released from polysomes upon stress is rapidly bound by proteins and forms ribonucleoprotein (RNP) condensates. Processing bodies (P-bodies) and stress granules (SG) are two types of RNP condensate that rapidly form within the cytoplasm in response to stress (Decker and Parker, 2012; Riggs et al., 2020). Despite differences in the RNA binding proteins involved, both P-bodies and SGs are formed through the condensation of translationally repressed mRNAs. When translation activity ceases, numerous P-bodies and SGs quickly emerge, exhibiting sizes ranging from 100 nm to microns in diameter, which can vary depending on the specific stress conditions and cell types (Jain et al., 2016; Rao and Parker, 2017). Polysomes and mRNA are extremely abundant, and therefore changes in the assembly state of ribosomes (from polysomes to monosomes) and the redistribution of mRNA to RNP condensates could potentially impact the biophysical properties of the cytoplasm.

Here, we investigated the impact of stress on mesoscale macromolecule diffusivity in the period immediately following acute glucose starvation in yeast. We discovered a transient increase in mesoscale cytoplasmic diffusivity that persists over the first hour of starvation. The collapse of polysome complexes was a prerequisite for this fluidization. In addition, the condensation of translationally arrested mRNAs dissociated into P-bodies in yeast or stress granules in mammalian cells was required for enhanced macromolecular diffusivity. This fluidization of the cytoplasm promoted the formation of Q-bodies, mesoscale structures that are essential for stress adaptation. Our findings provide new insights into how mRNA condensation influences the biophysical properties of cells, allowing for efficient reorganization of macromolecules.

## Results

### Mesoscale diffusivity transiently increases upon acute glucose starvation

Microrheology is the inference of the physical properties of a material from the observation of the motion of passive probes embedded within the environment (Mason and Weitz, 1995). Passive rheological probes have proven to be effective tools for inferring the biophysical properties of different cellular compartments (Chambers et al., 2022; Delarue et al., 2018; Munder et al., 2016; Parry et al., 2014; Plante et al., 2023; Shu et al., 2021; Xiang et al., 2021).

Previous reports have described decreased motion of macromolecules upon acute glucose starvation (Joyner et al., 2016). However, these observations monitored diffusivity of mRNP probes, which were later found to strongly interact with P-bodies, preventing the inference of general cytoplasmic properties from changes in their diffusivity (Heinrich et al., 2017). Furthermore, our recent study using passive rheological probes identified increased diffusivity at the mesoscale upon acute glucose starvation (Xie et al., 2023). To follow up on this observation, we sought to characterize the dynamics and mechanisms underlying this change. Therefore, we expanded our analysis to include additional time points. In addition, we investigated physical properties of the cytoplasm at two length-scales by using both µNS particles (50-150 nm diameter) (Parry et al., 2014) and 40nm-GEMs (40 nm diameter) (Delarue et al., 2018).

Note that the cytoplasm does not satisfy the simplifying assumptions of the Stokes-Einstein equation (Einstein, 1905; Stokes, 1901), therefore it is not possible to assign a single diffusion constant for a particle. Rather, the effective diffusivity of particles must be assigned at a specific time-scale. We chose a timescale of 100 ms for 40nm-GEMs and 1 second for µNS particles because the step-size distribution of these particles was similar to one another at these time intervals (Supplementary figure 1A). We denote the effective diffusion coefficient of 40nm-GEMs as D_eff_100ms_ and that of µNS particles as D_eff_1s_.

In agreement with our previous results (Xie et al., 2023), we found a transient increase in the D_eff_1s_ of μNS particles immediately after glucose starvation. This increased diffusivity of µNS was most significant at the first time point that we assayed; 15 minutes after acute glucose starvation μNS particles nearly doubled their D_eff_1s_ (Figure 1B). The step size distribution at 1 second also shifted to larger step sizes, confirming this result (Figure 1C). The anomalous exponent α value of μNS particles did not change significantly (Figure 1D). After 1-hour, the D_eff_1s_ of μNS particles returned to initial values (Figure 1B). Therefore, there is a significant fluidization of the cytoplasm at the ∼100 nm length-scale, but this physical change is transient, lasting less than one hour, perhaps explaining why it has not been reported previously.

40nm-GEMs also displayed a transient increase in D_eff_100ms_ and increased step sizes at 100 ms after 15 minutes of glucose starvation, but this was a mild effect compared to µNS at the 1 s timescale (Figures 1E and 1F). The anomalous exponent α value of 40nm-GEMs did not change significantly (Figure 1G). 40nm-GEMs D_eff_100ms_ returned to initial values at 30 minutes followed by a continuous decrease in diffusivity, reaching about 5-fold reduction at 120 min (Figure 1E). Both 40nm-GEMs and μNS particles displayed low diffusivity (less than 0.05 μm^2^/sec) after 2-hours of glucose starvation (Figures 1B and 1E). These observations suggest that the cytosolic environment changes in a dynamic way upon acute glucose starvation depending on length-scale. Larger particles (∼100 nm diameter) experience a transient reduction in constraint immediately after starvation. This effect, although less pronounced, is also apparent for smaller particles (40 nm diameter). At longer times, there is a reduction in diffusivity at both length-scales.

### ATP depletion does not cause increased diffusivity

Multiple factors could contribute to the increased diffusivity of μNS particles immediately after acute glucose starvation, and the subsequent decrease of diffusivity at longer times. One possible cytoplasmic alteration that might have widespread effects is the sudden reduction in intracellular ATP levels and pH upon glucose removal. Glucose is metabolized by glycolysis to produce ATP during proliferative growth. Budding yeast are adapted to acidic environments, and the plasma membrane proton transporter PMA1 uses large amounts of ATP to maintain a near neutral intracellular pH (Joyner et al., 2016; Martínez-Muñoz and Kane, 2017). Previous studies demonstrated that ATP levels are reduced by 70% and intracellular pH decreases to around 6 after 30 minutes of glucose starvation (Gutierrez et al., 2022; Joyner et al., 2016). Therefore, we determined the ATP and intracellular pH level again in our acute glucose starvation experiment. Using a ratiometric fluorescent ATP sensor, QUEEN (Takaine et al., 2019; Yaginuma et al., 2014), we found that acute glucose starvation led to ATP reduction within 10 minutes, and further decreased over time (Supplementary figures 1B, 1C). We also used the ratiometric pH sensor, pHluorin (Miesenböck et al., 1998), and confirmed that the intracellular pH of proliferative cells (pH = 7.5) was rapidly reduced to around 6 after 10 minutes of glucose starvation and fell to around 5.5 after 30 minutes of glucose starvation (Supplementary figures 1D, 1E).

To specifically test how macromolecular diffusivity was influenced by the low ATP and cytoplasmic acidification, we tracked the diffusion of both μNS particles and 40nm-GEMs after depletion of ATP by switching yeast cells to medium buffered at pH 5.5 without glucose, supplemented with 2-deoxyglucose (2-DG, an inhibitor of glycolysis) and antimycin A (an inhibitor of mitochondrial respiration.) This condition was previously shown to drastically reduce movement of endogenous particles as well as μNS particles (Munder et al., 2016). Consistent with those results, we found that both μNS particles and 40nm-GEMs displayed a reduction of α values, effective diffusion, and a shift to smaller step sizes after 30 minutes of ATP depletion (Supplementary figures 1F-H). These observations support previous conclusions that metabolic activity fluidized the cytoplasm (Parry et al., 2014).

Together, these results show that depletion of ATP and cytosolic acidification decrease mesoscale macromolecule diffusivity. These effects may explain the ultimate decrease in diffusivity more than an hour after glucose starvation. However, the initial increase of mesoscale diffusivity must be due to additional intracellular changes that counteract this effect in the short term. We sought to identify those factors.

### Disruption of polysomes is required for transiently increased mesoscale diffusivity

In the cytoplasm, ribosome concentration is a major determinant of mesoscale macromolecular diffusivity. Inhibition of TORC1 signaling reduces ribosome concentration leading to increased diffusivity of both μNS particles and 40nm-GEMs (Delarue et al., 2018). However, this change relies on autophagy and takes more than an hour. Therefore, we wondered if other aspects of ribosome physiology could explain the rapid fluidization of the cytoplasm upon acute glucose starvation.

In rapidly proliferating yeast cells, more than 80% of ribosomes are actively engaged in protein translation (Boehlke and Friesen, 1975; Waldron et al., 1977). Depending on the transcript length and translation efficiency, several ribosomes can simultaneously translate one mRNA, resulting in polysome complexes (polysomes) (Arava et al., 2003; Ingolia et al., 2009; Miller et al., 1970). Upon acute glucose starvation, various mechanisms rapidly inhibit translation initiation leading to disruption of polysomes (Janapala et al., 2019). Ribosomes dissociate from mRNA, disassembling into 40S and 60S ribosomal subunits (Ashe et al., 2000) and then rapidly reassemble into 80S monosomes in STM1-dependent process (Balagopal and Parker, 2011; Ben-Shem et al., 2011).We hypothesized that collapse of polysomes and assembly of monosomes might help fluidize the cytoplasm in stress conditions.

Consistent with previous studies (Ashe et al., 2000), we observed a dramatic reduction in polysomes upon acute glucose starvation (Figures 2A, 2B). Our hypothesis predicts that preventing polysome disassembly should prevent cytoplasmic fluidization. To test this prediction, we treated cells with cycloheximide. Cycloheximide freezes translation, prevents ribosome dissociation from mRNAs, and therefore prevents polysome collapse (Schneider-Poetsch et al., 2010) (Figures 2A, 2B). Consistent with our prediction, simultaneous glucose starvation and cycloheximide treatment completely abolished the initial increase in diffusivity of both μNS particles and 40nm-GEMs (Figures 2C, 2D). Instead, diffusivity of both particles immediately began to decrease. Cycloheximide treatment in glucose rich medium did not change the diffusivity of μNS particles or 40nm-GEMs (Supplementary figures 2A-D), suggesting that inhibition of translation *per se* does not greatly impact cytoplasmic fluidity, but rather changes in polysome structure are crucial. Therefore, we conclude that polysome disassembly is crucial for the initial transient increase of macromolecular diffusivity following glucose starvation (Figures 2C, 2D).

**Figure 2.**
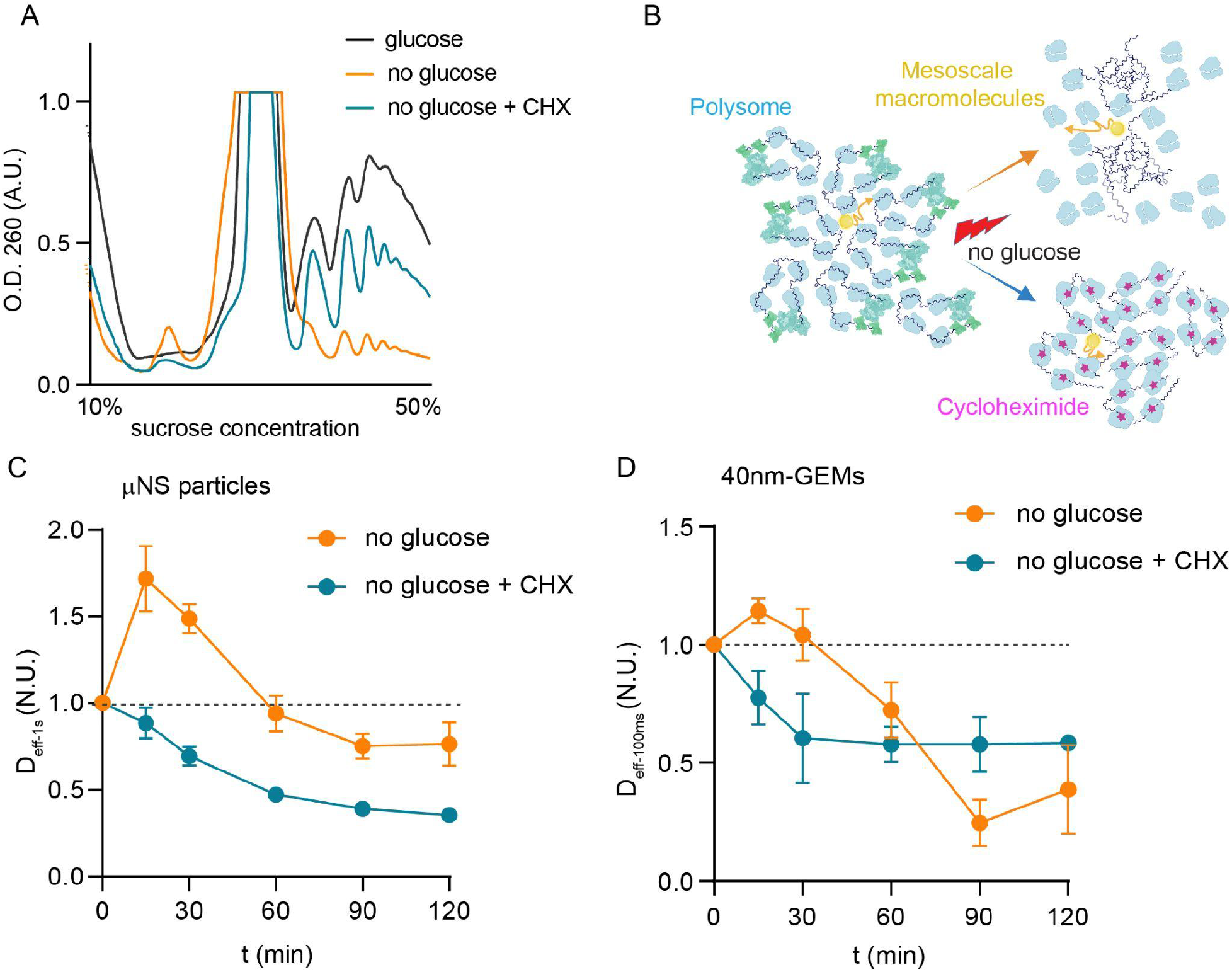
Polysome disassembly is necessary for increased mesoscale diffusivity in response to stress. **A.** Polysome profile analysis by fractionation of cell extracts on a sucrose gradient in the indicated conditions, cells were collected at 30 minutes upon switching to the respective medium (CHX: cycloheximide). **B.** Schematic of polysome rearrangement upon acute glucose starvation with or without 100 µg/ml cycloheximide. **C-D,** Fold change of median effective diffusion coefficients for μNS particles and 40nm-GEMs from 3 biological replicate experiments at the indicated conditions (CHX: cycloheximide, mean ± SEM.).

### Formation of P-bodies increases mesoscale diffusivity upon acute glucose starvation

Upon polysome disassembly mRNAs are suddenly released. These mRNA molecules quickly associate with RNA binding proteins that drive the formation of stress induced condensates, including processing bodies and stress granules (Buchan et al., 2008; Lui et al., 2010; Stoecklin and Kedersha, 2013) (Figure 3A). The release of mRNA and formation of these condensates is blocked by cycloheximide (Teixeira et al., 2005). We hypothesized that reorganization of mRNA into condensates might play an independent role in the rapid cytoplasmic fluidization by preventing the formation of local cages of disorganized mRNA immediately after glucose starvation.

**Figure 3.**
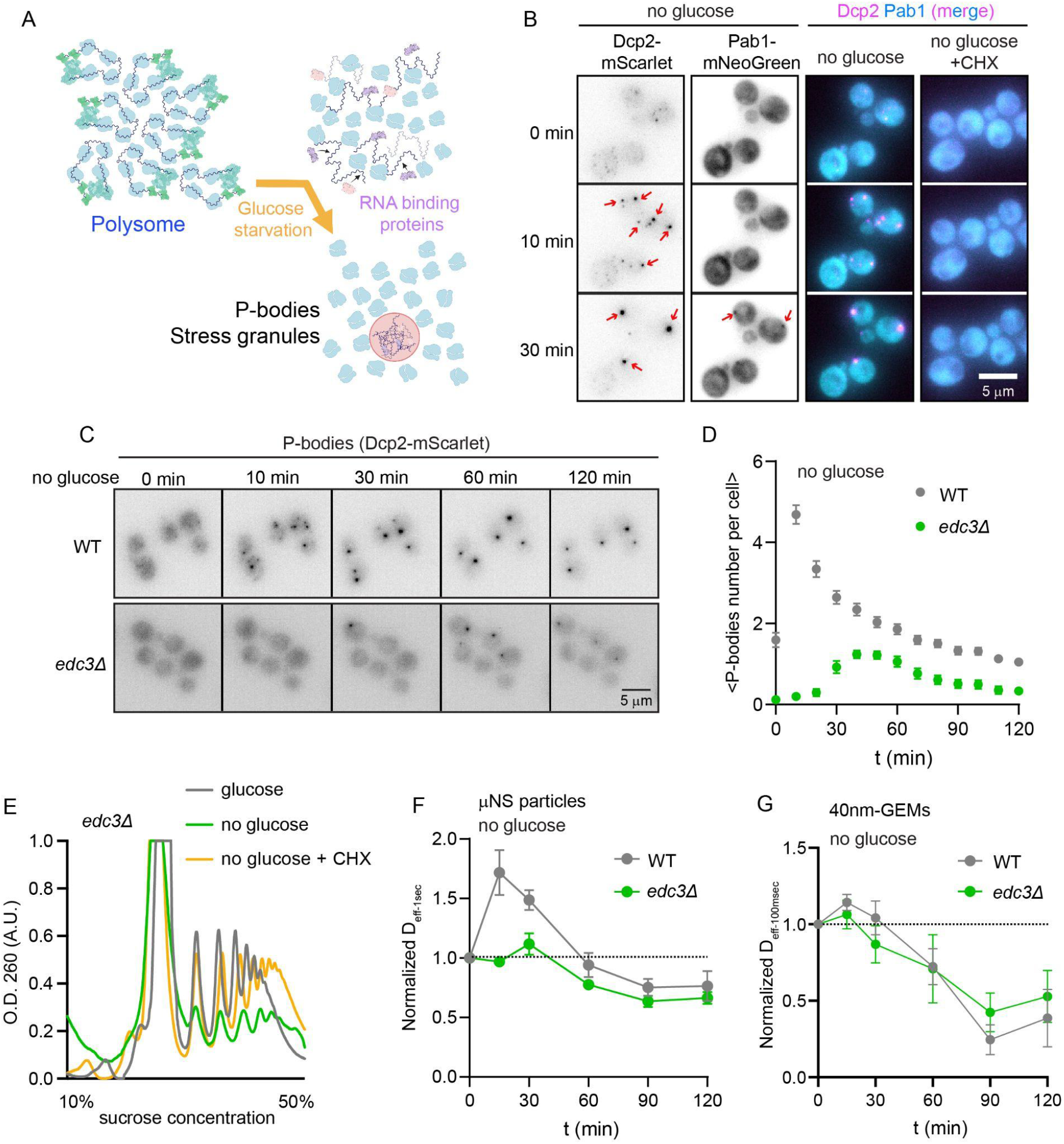
Formation of P-bodies increases the mesoscale particle diffusivity upon acute glucose starvation. **A.** Schematic of P-bodies and stress granules formation after glucose starvation. **B.** Representative fluorescent live cell images of P-bodies (Dcp2-mScarlet) and stress granules (Pab1-mNeoGreen) upon acute glucose starvation. Cycloheximide (CHX) treatment inhibits the formation of both RNP condensates. **C.** Time course showing formation of P-bodies (Dcp2-mScarlet marker) after acute glucose starvation in wild type and *edc3Δ S. cerevisiae* cells. **D.**Quantification of the number of Dcp2-mScarlet foci per cell at the indicated time points (wild type: n=64 cells; *edc3Δ*: n=51 cells, from 3 biological replicates). **E.** Polysome profile analysis by fractionation of *edc3Δ* cell extracts on a sucrose gradient in the indicated conditions, cells were collected at 30 min upon switching to the respective medium (CHX: cycloheximide). **F-G.** Fold change of median effective diffusion coefficients for μNS particles and 40nm-GEMs from 3 biological replicates in the indicated conditions (mean ± SEM.).

To test this hypothesis, we monitored the formation of RNP granules upon acute glucose starvation through live cell imaging of the Dcp2 and Pab1 proteins, which are markers for P-bodies and stress granules respectively (Figure 3B). Consistent with previous reports (Buchan et al., 2008), P-bodies formed rapidly, becoming clearly visible 10 minutes after acute glucose starvation, while stress granules appeared more slowly and were less prominent (Figure 3B). Treatment with cycloheximide completely inhibited the formation of both P-bodies and stress granules (Figure 3B).

Since the formation of P-bodies correlates with cytoplasmic fluidization, we tested if these structures are required for this physical change. The Edc3 protein is a key P-body scaffold (Currie et al., 2023; Decker et al., 2007; Xing et al., 2020), We found that an *EDC3* gene deletion mutant (*edc3Δ)* did not efficiently form P-bodies after glucose starvation, as indicated by a marked reduction in the appearance of Dcp2 foci (Figures 3C, 3D). We confirmed that polysomes were still disassembled after glucose starvation in *edc3Δ* cells (Figure 3E), consistent with previous reports (Decker et al., 2007). Based on these observations, we concluded that the reduction of microscopically visible P-bodies in an *edc3Δ* mutant was not due to defects in translational repression, but rather the absence of Edc3 prevented the condensation of untranslated mRNAs into P-bodies.

We tested if increased macromolecular diffusivity immediately after glucose starvation is attenuated in *edc3Δ* cells. First, we found neither rheological probe was sequestered into P-bodies upon acute glucose starvation (Supplementary figure 3A), making us confident that they were reliable reagents for defining the cytosolic environment outside of P-bodies. We observed almost no increase of μNS particle diffusivity in *edc3Δ* mutant cells, and also substantially decreased diffusivity of 40nm-GEMs during the initial hour of starvation (Figures 3F, 3G). Interestingly, at later time points (> 90 min), 40nm-GEMs diffusivity did not decrease as much in *edc3Δ* mutant cells as in wildtype cells. This effect was similar to the prevention of polysome disassembly, and indicated a less efficient late transition to a glassy cytoplasm. There was still a very slight initial increase in diffusivity of 40nm-GEMs and μNS particles in *edc3Δ* mutant cells. This increase was completely blocked when polysome disassembly was prevented by cycloheximide treatment (Supplementary Figures 3B-C). Together, these results indicate that conversion of polysomes to monosomes and condensation of mRNAs in P-bodies act synergistically to fluidize the cytoplasm upon glucose starvation.

### Increased mesoscale diffusivity facilitates Q-body formation upon acute glucose starvation

Multiple types of membraneless organelles form in stress conditions (Iserman et al., 2020; Kroschwald et al., 2018; Rao and Parker, 2017; Riggs et al., 2020; Sontag et al., 2022). In most cases, the formation of these structures occurs by an initial nucleation of small assemblies at multiple sites, and these then fuse together to form the mature organelle. This growth of condensates by coalescence can be inhibited by crowded or caged environments because subassemblies reach a size that can no longer diffuse efficiently to encounter one another and fuse together. For example, the growth of synthetic condensates is mechanically inhibited by chromatin (Lee et al., 2021). We therefore hypothesized that the fluidization of the cytoplasm by polysome disassembly and P-body formation could facilitate the formation of new membraneless organelles.

Q-bodies are membraneless organelles that are formed upon glucose starvation or heat shock. These stresses can lead to the formation of protein aggregates and proteotoxic stress (Sathyanarayanan et al., 2020; Takaine et al., 2022). Therefore, molecular chaperones act to facilitate compartmentalization of protein aggregates. One chaperone protein, Hsp42 has been demonstrated to efficiently associate with misfolded proteins and promote soluble Q-bodies formation. Deletion of the *HSP42* gene reduces Q-bodies assembly and cellular fitness when yeast cells are challenged by heat stress or glucose starvation (Escusa-Toret et al., 2013; Sathyanarayanan et al., 2020), indicating that Q-bodies help cells adapt to stress.

We tested if transiently increased mesoscale diffusivity upon glucose starvation was important for efficient Q-body assembly. We used endogenously tagged Hsp42-mScarlet to compare Q-body formation upon acute glucose starvation between wild type and *edc3Δ* mutant cells (Figure 4A). In wild type cells, we found that the average number of Hsp42-mScarlet foci increased over the first hour of glucose starvation until, on average, all cells had one Q-body (Figure 4B, black line.) Thus, Q-body formation is coincident with the transient fluidization of the cytoplasm after glucose starvation. Strikingly, Q-body formation was significantly attenuated in the *edc3Δ* mutant that does not effectively fluidize its cytoplasm (Figures 4A and 4B orange line) consistent with cytoplasmic fluidization enabling formation of mesoscale Q-bodies.

**Figure 4.**
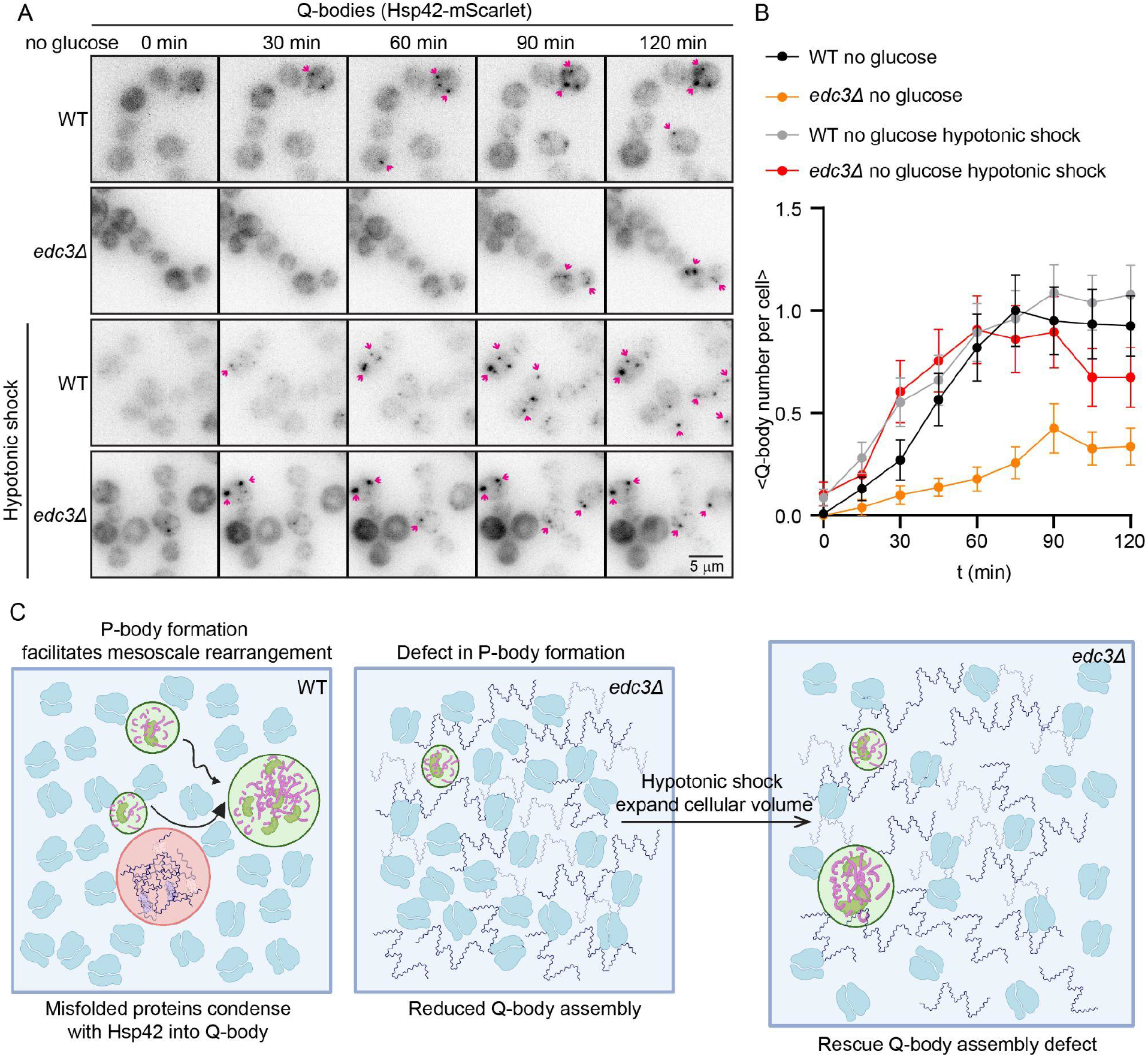
Increased mesoscale diffusivity is crucial for Q-body assembly upon acute glucose starvation. **A.** Representative fluorescent live cell images of Q-bodies, visualized by Hsp42-mScarlet, in wild type and *edc3Δ S. cerevisiae* cells upon acute glucose starvation. Hypotonic shock was achieved by overnight culture of yeast cells in synthetic complete media with 2% glucose and 500 mM KCl. Upon reaching log phase (O.D. 600nm < 0.4) cells were switched to starvation media of lower osmolarity (with no KCl.) **B.** Average number of Hsp42 foci per cell after acute glucose starvation in the indicated conditions (wild type -glucose: n=122 cells; *edc3Δ* -glucose: n=101 cells; wild type -glucose, hypotonic shock: n=103 cells; *edc3Δ* -glucose, hypotonic shock: n=86 cells, N=3 biological replicates on independent days, mean ± SEM.). **C.** Graphic illustration of Q-bodies assembly, created with BioRender.

We sought to distinguish whether P-bodies facilitate Q-body formation through fluidization of the cytoplasm or through an alternative mechanism. If physical fluidization of the cytoplasm is the key factor, we expect to be able to rescue Q-body formation by fluidizing the cytoplasm of *edc3Δ* mutant cells using methods that do not require P-body formation. It is possible to decrowd the cytoplasm using acute hypotonic shock (Delarue et al., 2018). To achieve hypotonic shock, we cultured both wild type and *edc3Δ* mutant yeast cells in hypertonic media (media supplemented with 500 mM KCl as an osmolyte) overnight. In these conditions, cells activate the high-osmolarity glycerol (HOG) pathway to accumulate osmolytes (primarily glycerol), thereby osmotically balancing the cell interior enabling growth (Brewster et al., 1993). Suddenly shifting cells to media without excess osmolyte (without KCl) creates a hypotonic shock because the osmolytes that have accumulated in the cell take time to be degraded and exported (Hohmann et al., 2007). This hypotonic shock causes water to enter the cell, leading to a sudden increase in volume and a corresponding decrease in macromolecular crowding, thereby increasing mesoscale diffusivity (Delarue et al., 2018).

Upon hypotonic shock, the diffusivity of μNS particles increased drastically at 5 minutes in both wild type and *edc3Δ* mutant cells, without changing the overall μNS particle size (Supplementary figures 4A-C). Experimentally-induced decrowding of the cytoplasm was sufficient to completely rescue Q-body assembly in *edc3Δ* cells (Figures 4A-C, red line). These results support our hypothesis that P-body formation increases mesoscale macromolecular diffusivity thereby enabling the formation of mesoscale Q-body condensates during the initial response to glucose starvation.

### Stress granule formation increases mesoscale diffusivity in mammalian cells

Our results demonstrate that stress induced condensation of mRNA into P-bodies increases mesoscale diffusivity in yeast cells. We next wondered how general the physical effect of mRNA condensation might be, and if there are similar consequences for stress-induced RNP formation in the mammalian cytoplasm. We initially investigated P-bodies formation in U2OS mammalian cells, but found that P-bodies are constitutively present, independent of stress, and so not amenable to addressing our question (Supplementary figure 5A). Therefore, we focused on stress granule formation in mammalian cells.

Stress granules are induced upon many types of stress and are widely implicated in various physiological or pathological conditions in mammalian cells (Ash et al., 2014; Guillén-Boixet et al., 2020; Lavalée et al., 2021; Riggs et al., 2020; Yang et al., 2020). We sought to test whether formation of stress granules upon translational arrest changes the physical properties of the mammalian cell cytoplasm. We initially tried external stresses to induce stress granule formation but found that the typical inducing agents (oxidative stress, heat shock, or osmotic compression) had pleiotropic effects or caused physical changes by independent mechanisms that were difficult to deconvolve. However, translation inhibition is sufficient to drive stress granule formation. Therefore, we used translation inhibitors to trigger the formation of stress granules in mammalian cells (Figure 5A).

**Figure 5.**
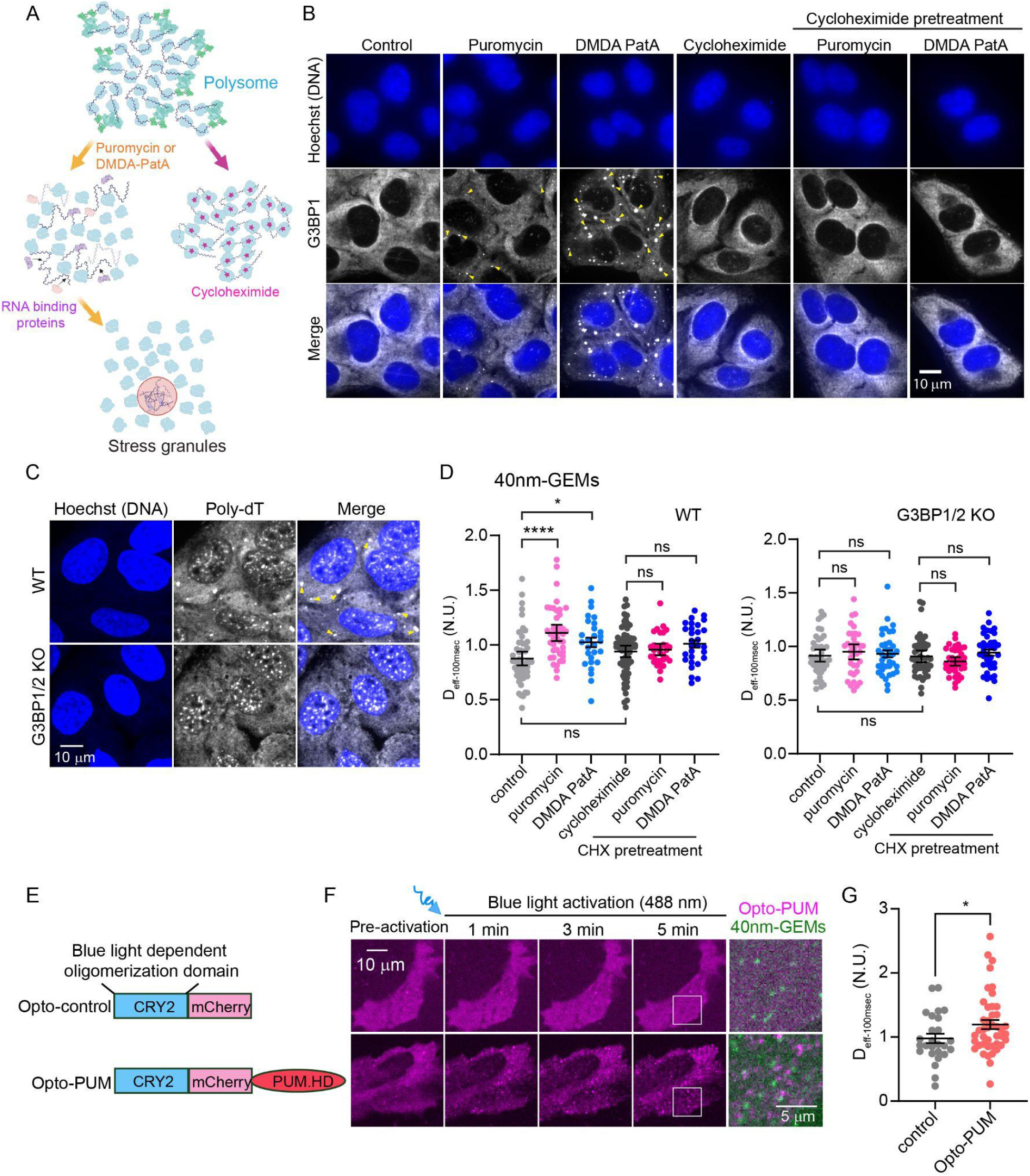
Partitioning of mRNAs into RNP granules increases mesoscale macromolecular diffusivity in mammalian cells. **A.** Graphic illustration of the mode of action of various translational inhibitors, and their predicted relationship to stress granule formation (created with BioRender). **B.** Representative immunofluorescence images of stress granules (G3BP1) in U2OS cells treated with indicated protein translation inhibitors for 30 minutes. Hoechst dye was used to stain DNA. **C.** Representative poly-dT FISH images to analyze the physical distribution of poly-adenylated mRNAs in wild type and G3BP1/2 knockout (KO) U2OS cells after 10 nM DMDA-PatA treatment for 30 minutes. Hoechst dye was used to stain DNA. **D.** Fold change of median effective diffusion coefficients (D_eff-100ms_) for 40nm-GEMs in the indicated conditions. The median D_eff-100ms_ was measured for the same cell before and 30 minutes after each treatment. Each dot represents one cell, the value plotted as the D_eff-100ms_ in the treatment condition normalized to the D_eff-100ms_ for the same cell prior to treatment (0 minutes). (Three independent experiments were performed with individual cell measurements represented by each dot, WT control n=57; WT puromycin n=42; WT DMDA-PatA n=28; WT cycloheximide n=67; WT Cycloheximide pretreatment puromycin n=31; WT Cycloheximide pretreatment DMDA-PatA n=29. G3BP1/2 KO control n=44; G3BP1/2 KO puromycin n=36; G3BP1/2 KO DMDA-PatA n=37; G3BP1/2 KO cycloheximide n=46; G3BP1/2 KO Cycloheximide pretreatment puromycin n=38; G3BP1/2 KO Cycloheximide pretreatment DMDA-PatA n=38. mean ± SEM. Statistical comparison was by a one-way ANOVA test. ns=not significant, *p<0.05, **p<0.01, ****p<0.0001). **E.** Graphic illustration of blue light inducible Opto-control and Opto-PUM constructs. **F.** Representative live cell images of U2OS cells expressing Opto-Control or Opto-PUM together with 40nm-GEMs, 488 nm blue light laser was stimulated every 5 seconds for a period of 5 minutes to drive Opto-PUM condensation. **G.** Fold change of median effective diffusion coefficients (D_eff-100ms_) for 40nm-GEMs from 3 biological replicate experiments in the indicated conditions (The same cells D_eff-100ms_ at 30 min was normalized by D_eff-100ms_ at 0 min, n=26 cells for Opto-control, n=43 for Opto-PUM, mean ± SEM. Statistical comparison was performed using a paired two-tailed t test. *p<0.05.).

Translation inhibitors that cause collapse of polysomes and release of mRNA (puromycin, which causes premature chain termination during translation, and DMDA-PatA, which inhibits the translation initiation factor eIF4A) (Baliga et al., 1970; Dang et al., 2006; Kommaraju et al., 2020; Kudla and Karginov, 2016), induced widespread formation of stress granules as indicated by condensation of the stress granule protein G3BP1 (Figure 5B). In contrast, treatment with the translation inhibitor cycloheximide, which stalls translation and prevents polysome disassembly and the release of untranslated mRNAs (Schneider-Poetsch et al., 2010), did not lead to stress granules formation (Figure 5B). Moreover, pretreatment of cycloheximide before applying puromycin or DMDA PatA suppressed stress granule formation (Figure 5B), supporting the idea that release of mRNAs from polysomes seeded the formation of stress granules.

We also analyzed the overall spatial distribution of mRNAs in the cell using poly-dT Fluorescence In Situ Hybridization (FISH) staining after triggering stress granule formation using DMDA-PatA. In U2OS wild type cells, we observed clear foci of concentrated mRNA in the cytoplasm after DMDA-PatA treatment, consistent with mRNA condensation into stress granules (Figure 5C).

G3BP1 and G3BP2 proteins interact with both RNAs and RNA binding proteins and are crucial scaffolds for stress granules formation (Guillén-Boixet et al., 2020; Yang et al., 2020). Double mutant G3BP1/2 knockout cells were previously demonstrated to prevent stress granule formation in some types of stress such as sodium arsenite (Yang et al., 2020). If stress granules were responsible for the condensation of mRNA observed by FISH, then mRNA foci should not form in G3BP1/2 knockout cells. Consistent with this prediction, mRNA clusters were absent in G3BP1/2 knockout cells (Figure 5C and supplementary figures 5B-C.) Together, these results support the model that mRNA released from polysomes upon translation inhibition condenses into stress granules in mammalian cells.

We also observed an increased number of P-bodies in response to translational inhibition as indicated by immunofluorescence staining of Dcp1 (Figure 5C and supplementary figures 5D-E.) However, P-bodies were also present in control conditions, therefore we did not focus on these condensates.

We predicted that mRNA condensation into stress granules could change the biophysical properties of the cytoplasm. To test this hypothesis we expressed cytosolic 40nm-GEMs in wild type and G3BP1/2 knockout cells to measure mesoscale macromolecule diffusivity. Note that μNS particles were originally derived from a vertebrate viral protein (Broering, 2002), and so we were not confident that these could be used as orthogonal rheological probes. Interestingly, upon puromycin and DMDA-PatA treatment, 40nm-GEMs showed an increase of diffusivity in wild type U2OS cells but not in G3BP1/2 knockout cells (Figure 5D). Cycloheximide pretreatment of cells suppressed this effect when cells were treated with either puromycin or DMDA-PatA (Figure 5D). Therefore, conditions that lead to formation of stress granules cause increased mesoscale macromolecular diffusivity, and treatments that prevent stress granule formation abolish this physical change.

To test whether mRNA condensation, independent of translation inhibition and polysome collapse, fluidizes the mammalian cell cytoplasm, we developed a strategy to induce synthetic mRNA condensates without cellular stress. Artificially assembled RNP granules containing the pumilio homology domain (PUM.HD) from the human Pumilio 1 protein were previously demonstrated to condense mRNA (Cochard et al., 2022; Garcia-Jove Navarro et al., 2019). We modified the artificial PUM.HD-RNP system by adding the homo-oligomerization cryptochrome 2 (CRY2) domain from *Arabidopsis thaliana (Bugaj et al., 2013; Che et al., 2015)*, so that we could use blue light to control its dynamic assembly. We named this light-inducible condensate “Opto-PUM” (Figure 5E) We also made a CRY2-mCherry construct with no RNA binding domain, and named this “Opto-control” (Figure 5E). Within minutes of blue light activation, we observed the formation of numerous Opto-PUM foci, while no foci were formed in Opto-control cells (Figure 5F). We coexpressed 40nm-GEMs to quantify mesoscale diffusivity. 40nm-GEMs did not partition into Opto-PUM foci, indicating that they can be used to infer diffusivity (Figure 5F). We observed a significant increase in the diffusivity of 40nm-GEMs upon formation of Opto-PUM foci (Figure 5G.) This result is consistent with the model that mRNA condensation is sufficient to fluidize the cytoplasm, independent of translation inhibition associated with cellular stress.

## Discussion

The physical properties of the cytoplasm are likely to profoundly impact a broad range of biochemistry. Growing evidence shows that these properties can dramatically change, especially in stress conditions, but the mechanisms for these changes and their biological consequences remain poorly understood. We have discovered a dynamic physical change in the cytoplasm that leads to a transient increase in mesoscale diffusivity in response to acute stress. We demonstrate that this enhanced fluidization of the cytoplasm is important for the formation of Q-bodies, stress-induced structures that are required for sequestration and clearance of misfolded proteins that accumulate as a result of stress.

Ribosome density was previously shown to be a key determinant of cytoplasmic macromolecular crowding (Delarue et al., 2018). However, the role of higher-order ribosome organization was not addressed in that study. Here, we found that assembly of ribosomes and mRNAs into polysomes is a key determinant of mesoscale motion. We propose that polysomes form cages in the cytoplasm that hinder mesoscale diffusion. This could be due to their large size, interactions between mRNA molecules within the polysomes, or RNA-binding proteins involved in translation; the exact physical interactions still require elucidation. However, it is clear from our cycloheximide experiments that polysome disassembly is required for cytoplasmic fluidization. The importance of higher order ribosome organization for the physical properties of the cytoplasm provides a general mechanism for rapid regulation as almost all stresses lead to translational arrest and disassembly of polysomes.

Polysome disassembly is necessary but not sufficient for cytoplasmic fluidization; rather, we found that mRNA condensation was essential. Without this condensation, we propose that mRNA forms networks in the cytoplasm that create local cages. In yeast, the crucial condensates were P-bodies. In mammalian cells, P-bodies were constitutively present in the cytoplasm precluding the study of their dynamics. This could be due to the non-physiological environment of the imaging dish, or the use in our study of cancer cells that may have adapted their physical properties during oncogenesis. However, we found that stress granule formation led to cytoplasmic fluidization upon translational inhibition in mammalian cells and that deletion of the key stress granule scaffolds G3BP1 and G3BP2 was sufficient to prevent the physical change. It will be interesting to further characterize the biophysical roles of P-bodies in mammalian cells. We speculate that different mRNA condensates may be more important in different contexts and stresses, or that evolutionary divergence has altered the contribution of stress granules and P-bodies in yeast and mammalian cells. Despite this difference, the fundamental processes of polysome collapse and mRNA condensation into mRNP granules is an evolutionarily conserved mechanism for stress-induced fluidization of the cytoplasm (Figure 6).

**Figure 6.**
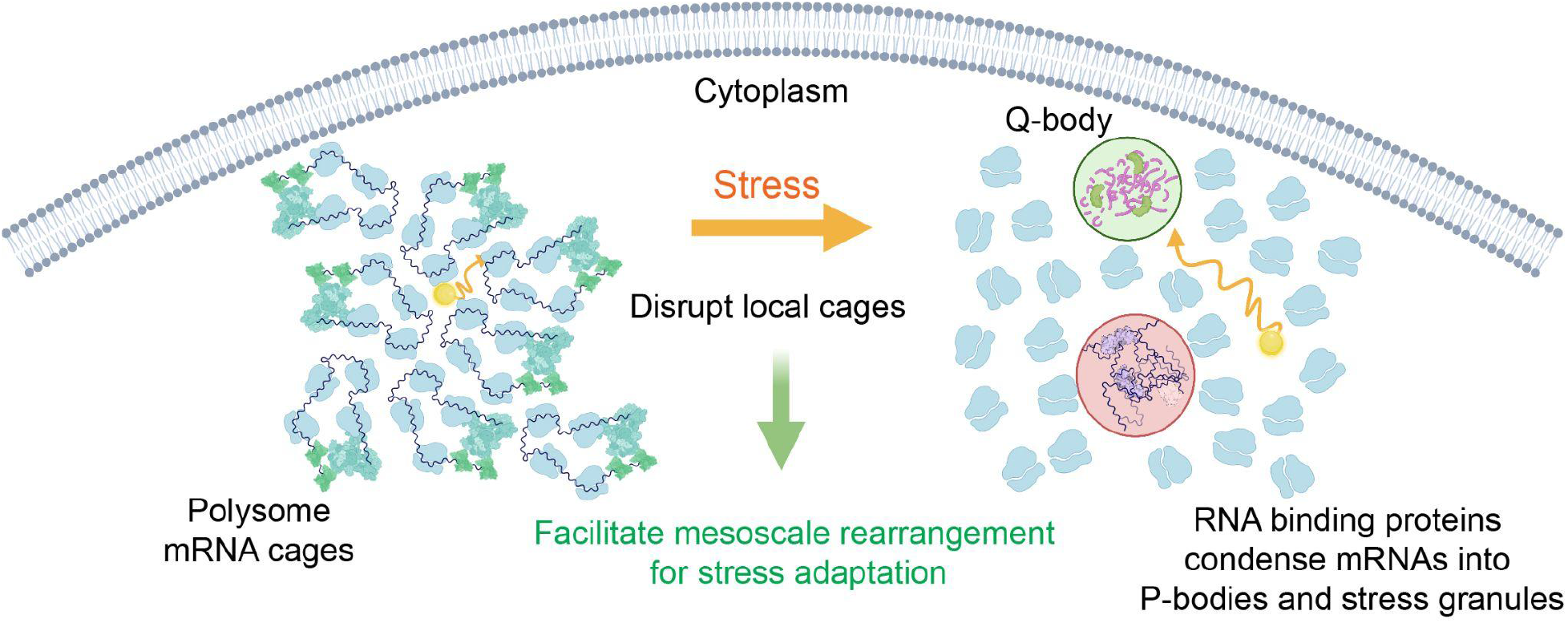
Polysome collapse and mRNP granule formation transiently fluidize the cytoplasm enabling effective stress response. Upon stress-induced translational arrest, the disassembly of polysomes results in the release of untranslated mRNAs. These mRNAs condense through the action of RNA binding proteins, leading to the formation of processing bodies and stress granules. Both polysome collapse and mRNA condensation are required for increased mesoscale diffusivity. This fluidization of the cytoplasm enables the assembly of mesoscale structures such as Q-bodies that are required for stress adaptation.

Several studies have demonstrated that decreases in ATP cause yeast or bacterial cells to transition to a glassy/solid state in the cytoplasm (Munder et al., 2016; Parry et al., 2014; Plante et al., 2023). In our study, we also found that the reduction of ATP and cytoplasmic pH eventually results in decreased mesoscale diffusivity. However, this effect is initially counteracted by polysome collapse and partitioning of untranslated mRNAs into P-bodies leading to fluidization of the cytoplasm during the immediate response to starvation. At later times, mesoscale diffusivity decreases to levels that were described previously, ultimately resulting in a glassy cytoplasm. In fact, the disassembly of polysomes and condensation of untranslated mRNAs into P-bodies appears to facilitate the ultimate glassy/solid state transition.

Preventing polysome disassembly with cycloheximide or suppression of P-body formation through loss of Edc3 prevented transient fluidization, but these conditions also resulted in higher diffusivity at later time points, at least at the 40 nm length-scale (Figures 2D, 3G and supplementary figure 3C) Therefore, mRNA condensation and conversion of polysomes to monosomes may also be beneficial for the eventual transition to a solid-like state that has been proposed to facilitate dormancy (Munder et al., 2016).

The dynamic biophysical changes are likely to have widespread impacts on biochemistry. We propose that the initial fluidization of the cytoplasm enables rapid reorganization of the cell enabling adaptation to the stress condition. In all stresses, proteins tend to misfold, and the creation of mesoscale membraneless organelles is an important mechanism to allow for refolding or degradation (Escusa-Toret et al., 2013). Phase separation of membraneless organelles requires mesoscale subassemblies to fuse together, a process that can be inhibited by mesoscale caging effects (Lee et al., 2021). We found that the initial fluidization of the cytoplasm was important for the formation of Q-bodies. It will be interesting to investigate if other mesoscale assemblies that are known to assemble during ATP depletion (Marini et al., 2020; Munder et al., 2016; Plante et al., 2023) also require cytoplasmic fluidization for their formation.

In addition to the formation of protein aggregates, changes in membrane structure have been observed during glucose starvation in yeast. Endocytosis and exocytosis change the complement of carbon transporters (Feyder et al., 2015), a process that requires efficient trafficking of 100 nm vesicles. There are also large scale changes in organelle structure, including mitochondrial morphology changes, (Laporte et al., 2018), changes in inter-organelle connectivity, including mitochondria-ER contacts, and the nuclear-vacuole junction (Wood et al., 2020). These findings indicate that membrane reorganization within the cytoplasm is critical for ensuring cell survival during long-term starvation. Therefore, it will be interesting to investigate whether the fluidization of the cytoplasm also facilitates reorganization of membrane-bound organelles.

We have demonstrated that the formation of P-bodies in yeast cells enables the efficient compartmentalization of misfolded proteins into Q-bodies during stress adaptation. It will be interesting to further explore the possible biophysical and functional roles of stress granule formation in mammalian cells. There is growing evidence for the involvement of stress granules in human diseases. For example, cancer cells with oncogenic KRAS mutations upregulate stress granules through the production of bioactive lipid prostaglandins, leading to the inhibition of translation initiation factor eIF4A. Enhanced stress granule formation in this context has been shown to increase tumor fitness (Grabocka and Bar-Sagi, 2016). These observations imply that stress granule formation confers a selective advantage to cancer cells. It will be interesting to investigate if biophysical effects of stress granule assembly contribute to this adaptation.

On the other hand, aberrant stress granule formation appears to be cytotoxic in neurodegenerative diseases (Zhang et al., 2019). The cytoplasmic pathology observed in end-stage neurodegeneration often involves the accumulation of unconventional stress granules containing mislocalized RNA binding proteins, such as aggregated FUS (Deng et al., 2014; Dormann et al., 2010) or mutated TDP43 with abnormal phosphorylation (Neumann et al., 2009). Misregulation of RNA binding proteins is likely to alter the assembly kinetics of stress granules, potentially impacting the biophysical properties of the cytoplasm. Our study motivates the detailed study of these properties in diseased neurons: aberrant crowding and fluidity could contribute to the proteostatic collapse that underpins all neurodegeneration.

P bodies were first identified more than two decades ago but their functional significance has remained opaque. Our findings provide key functional insights into their essential role. Furthermore, we reveal a unifying biophysical consequence of mRNA condensation into both P bodies and stress granules establishing a new framework to investigate the role of mRNA condensates in both normal physiology and disease.

## Methods

| Reagent or Resource | Source | Identifier |
| --- | --- | --- |
| **Antibodies** |  |  |
| Anti-G3BP1 | Santa Cruz | sc-81940 |
| Anti-Dcp1 | Abcam | ab183709 |
| Anti-Rabbit IgG (H+L) | Invitrogen | A11036 |
| Anti-Mouse IgG (H+L) | Invitrogen | A21236 |
| **Chemicals** |  |  |
| 2-Deoxy-D-glucose | Sigma-Alrich | D8375 |
| Antimycin A | Sigma-Alrich | A8674 |
| Cycloheximide | Sigma-Alrich | C7698 |
| puromycin | MedChem Express | HY-B1743A |
| des-methyl, des-amino pateamine A | Daniel Romo lab |  |
| **Experimental Models: Organisms/Strains** |  |  |
| *MATa his21 leu20 met150 ura30*<br>pINO4::pINO4-PFV-Sapphire-LEU2, | Holt lab | LH4248 |
| *MATa his21 leu20 met150 ura30*<br>pHIS3::pHis3-GFP-uNS::Ura3 | Holt lab | LH4544 |
| *MATa his21 leu20 met150 ura30* His3::pTEF -QUEEN | Holt lab | LH4819 |
| -2m-TCYC1 |  |  |
| *MATa his21 leu20 met150 ura30* Leu2::pTDH3-pHluorin-tTDH3 | Holt lab | LH4820 |
| *MATa his21 leu20 met150 ura30* Dcp2-mScarlet::HIS3, Pab1-mNeoGreen::NatNT | Holt lab | LH4821 |
| *MATalpha his21 leu20 met150 ura30* Dcp2-mScarlet::HIS3, Pab1-mNeoGreen::NatNT *edc3Δ::hphMX* | Holt lab | LH4822 |
| *MATa his21 leu20 met150 ura30* pHis3-GFP-uNS::Ura3, Dcp2-mScarlet::HIS3 | Holt lab | LH4823 |
| *MATa his21 leu20 met150 ura30* pINO4::pINO4-PFV-Sapphire-LEU2, Dcp2-mScarlet::HIS3 | Holt lab | LH4824 |
| *MATalpha his21 leu20 met150 ura30 edc3Δ::hphMX* pHis3-GFP-uNS::Ura3 | Holt lab | LH4560 |
| *MATa his21 leu20 met150 ura30 edc3Δ::NatRMX*, pINO4::pINO4-Pfv-Sapphire-LEU2 | Holt lab | LH4825 |
| *MATalpha his21 leu20 met150 ura30* Hsp42-mScarlet::HIS3 | Holt lab | LH4565 |
| *MATalpha his21 leu20 met150 ura30 edc3Δ::hphMX* Hsp42-mScarlet::HIS3 | Holt lab | LH4566 |
| HEK293T mammalian cell line | Holt lab |  |
| U2OS mammalian cell line | J. Paul Taylor lab |  |
| U2OS G3BP1/2 KO mammalian cell line | J. Paul Taylor lab |  |
| U2OS 40nm-GEMs mammalian cell line | Holt lab |  |
| U2OS G3BP1/2 KO 40nm-GEMs mammalian cell line | Holt lab |  |
| **Recombinant DNA** |  |  |
| Yeast plasmid pRS305-Leu2-pINO4-PFV-Sapphire | Holt lab | pLH497 |
| Yeast plasmid pRS306-pHIS3-GFP-μNS | Holt lab | pLH1125 |
| Yeast plasmid pFA6a-ymScarletI-SpHIS5 | Holt lab | pLH1688 |
| Yeast plasmid pFA6a-mNeonGreen-natNT2 | Holt lab | pLH1532 |
| Yeast plasmid pFA6a-hphMX6 | Holt lab | pLH1493 |
| Yeast plasmid pFA6a-natMX6 | Holt lab | pLH1494 |
| Yeast plasmid pTEF1-QUEEN-2m-tCYC1 | Masakatsu Takaine lab | pLH2156 |
| Yeast plasmid pRS305-pTDH3-pHluorin-tTDH3 | Holt lab | pLH1097 |
| Mammalian plasmid pUBC-PFV-Sapphire | Holt lab | pLH1984 |
| Mammalian plasmid pMD2.G | Addgene #12259 | pLH1863 |
| Mammalian plasmid psPAX2 | Addgene #12260 | pLH1864 |
| Mammalian plasmid pHR-mCherry-Cry2WT | Addgene #101221 | pLH1972 |
| Mammalian plasmid pHR-PUM.HD-mCherry-Cry2WT | Holt lab | pLH2126 |
| **Software** |  |  |
| NIS Elements | Nikon |  |
| FIJI | https://doi.org/10.1038/nmeth.2019 |  |
| Prism 9 | GraphPad |  |
| Python | https://www.python.org/downloads/release/python-370/ |  |

### Yeast strain construction

Endogenous protein tagging at the C terminal with fluorescent protein was constructed via transformation of PCR products containing fluorescent proteins with auxotrophic marker gene, using either plasmid pLH1688 or pLH1532. The PCR products contained 42bp of homology to the 5′ and 3′ of the target gene stop codon region. Deletion strains were constructed via transformation with PCR products containing antibiotic resistance cassettes (pLH1493 and pLH1494). PCR products contained 42bp of homology to the 5′ and 3′ genomic regions immediately adjacent to the gene to be deleted. To integrate either 40nm-GEMs or uNS particles into the yeast strain, plasmid pLH497 was linearized by restriction enzyme SnaBI, and pLH1125 was linearized by primers located at the HIS3 promoter region. To integrate the ATP sensor QUEEN or pH sensor pHluorin, plasmid pLH2156 (a gift from Satoshi Yoshida lab) was linearized by restriction enzyme PstI, plasmid pLH1097 was linearized by restriction enzyme KasI. All DNA products were transformed to the yeast strains based on LiAc approach according to standard protocol.

### Lentivirus production and cell transduction

HEK293T cells (9×10^6^ per 15 cm dish) were plated in antibiotic free DMEM (Gibco, Cat. No. 11995073) supplemented with 10%FBS (Gemini bio-products, Cat. no. 100-106). The next day, cells were transfected with transgene plasmid (pLH1984) together with lentivirus packaging plasmids psPAX2 and pMD2.G, using fuGENE HD™ transfection reagent (Promega, Cat. no. E2312) following manufacturer’s protocol. 24 hours later, 15 ml antibiotic free DMEM was replaced and the supernatants were collected at both 48 and 72 h post-transfection, and stored at 4°C. Virus titers were concentrated by centrifugation at 4,000 rcf for 40 minutes in an Amicon Ultra-15 30 KDa centrifugal filter (MilliporeSigma, Cat. No. UFC903024). Concentrated viral suspensions were aliquoted and stored at −80°C until later use. Lentivirus was introduced into U2OS wild type and G3BP1/2 KO cell lines of interest via reverse transduction with 1-10 μL of concentrated virus in fresh media, and replacing media after 24 hours. After cell lines stabilized, they were frozen in 10% DMSO (Sigma-Aldrich, Cat. no. D2650-100) in FBS (Gemini bio-products, Cat. no. 100-106) and thawed for use in experiments whenever needed.

### Mammalian cell culture and transient transfection

U2OS cells (Kindly provided by J. Paul Taylor lab) were cultured in DMEM containing high glucose and sodium pyruvate supplemented with 10%FBS and 50 U/ml penicillin, and 50 μg/ml streptomycin (Gibco™, Cat. No. 15140122), and maintained at 37°C in a humidified incubator with 5% CO_2_. Cells were regularly split in fresh medium upon reaching 80-90% confluency. All cells were routinely tested for mycoplasma by PCR screening of the conditioned medium.

For transfection, U2OS cells were seeded as 60-70% confluency in a 6-well glass bottom plate (Cellvis, Cat. No. P06-1.5H-N) on the day before transfection and were transfected with 1 μg of plasmid DNA per well (pLH1972 or pLH2126) using FuGENE HD™ reagent per manufacturer guidelines. 24 hour post-transfection, fresh DMEM medium was replaced. And imaging experiments were usually carried out between 24 to 48-hour post-transfection.

### Yeast cell culture for imaging

*Saccharomyces cerevisiae* strains were revived from -80°C freezer on YPD plate for overnight growth. The next day, a patch of yeast cells were inoculated into 5-mL synthetic complete media with 2% glucose (SCD), and the cultures were grown at 30°C in a rotating incubator for 5-6 hours without exceeding an O.D.600 of 0.4. Afterwards, the cultures were prepared as a few tubes with 4-6 times of 10x dilution, and culture for an overnight growth in order to reach O.D.600 between 0.1-0.4 for the next day ‘s imaging experiment. To perform acute glucose starvation, 96-well (Cellvis, Cat. no. P96-1.5H-Nor) or 384-well (Cellvis, Cat. no. P384-1.5H-Nor) glass bottom imaging plates were precoated with 1mg/ml concanavalin A (Alfa Aesar, Cat. no. J61221) before applying the yeast cell culture. 10 min later when cells settled down to the bottom of the imaging plate, the culture medium was removed completely, and four additional wash of cells with SCD medium, and then perform imaging recorded as the initial 0 min time point. Afterwards, SCD medium was removed, followed by four additional wash of the cell with treatment medium. Cells were then imaged at the correspondent time points afterwards.

To perform acute glucose, SC medium supplemented with 2% sorbitol (to balance the osmotic pressure exerted by 2% glucose) was used. To perform ATP depletion experiment, SC medium supplemented with 2% sorbitol, pH adjusted as 5.5, 2-Deoxy-d-glucose and antimycin A were used in combination at 20 mM and 10 μM respectively.

### Drug treatments

Cycloheximide was used at a final concentration of 100 μg/ml in all yeast experiments, while for mammalian cell experiments, 20 μg/ml of cycloheximide was used as a final concentration. Puromycin and des-methyl, des-amino pateamine A (DMDA PatA, a gift from Daniel Romo lab) were used at a final concentration of 100 μg/ml and 10 nM respectively in all mammalian cell culture experiments.

### Sucrose gradient to fractionate polysomes

60 mL yeast cell culture in SCD medium reaching O.D.600 around 0.4 were collected by 3000 g centrifugation. And cells were washed with treatment medium before switching to the correspondent medium for additional 30 min culture, and all cells were collected again by centrifugation and freshly frozen in -80°C freezer. To prepare sucrose gradient, four layers of sucrose solutions (2.7 ml each) were prepared in concentrations as 10%, 23.3%, 36.6% and 50%, where the sucrose were layered from bottom to top as high to low concentrations in 14×89 mm ultracentrifuge tubes (Beckman, 14×89 mm polypropylene centrifuge tubes, Cat. no. 331372), and each layer of sucrose was freshly frozen in -80°C for 20 min before adding a second layer on top. To prepare cell lysis, polysome buffer (10 mM Tris, 10 mM MgCl_2_, 100 mM KCl, 6 mM β-mercaptoethanol, 100 μg/ml cycloheximide, protease inhibitors (Thermo Scientific™, Cat. no. A32961) pH 7.5) were freshly prepared, and cell pellets were washed and then resuspended in 500 μl polysome buffer in the glass tube (Fisher Scientific, Cat. No.14-961-32). Afterwards, 0.5 mm glass beads (BioSpec Products, Cat. No.11079105) were added as half of the volume in cell suspension and vortexed four times as 30 sec each, with 1 min rest time on ice in between. And then all the cell lysis extract were transferred to a new 1.5 ml tube, and 300 ul polysome buffer were used to wash the glass beads then combined with the cell lysis extract. To clarify the cell lysis extract, 10 min of centrifugation at 15000 g was performed at 4°C. And the supernatant of cell lysis extract was transferred to a new 1.5 ml tube, where the OD260 was measured by spectrophotometer. 9 units of OD260 cell lysis extract of each sample was then layered on top of the sucrose gradient, all the samples were then centrifuged at 39,000 rpm (Beckman, SW 41 Ti Swinging-Bucket Rotor) at 4°;C for 2.5 hrs. Afterwards, the samples were subjected to the fractionation system for analysis (Brandel, SYN-202 density gradient fractionation system).

### Passive rheological probes imaging and single particle tracking

To image 40nm-GEMs in *Saccharomyces cerevisiae*, TIRF Nikon TI Eclipse microscope in highly inclined thin illumination mode (HILO) was used at GFP laser (49002-ET-EGFP) excitation with 100% power. Fluorescence was recorded with a scMOS camera (Zyla, Andor) with a 100x Phase, Nikon, oil NA = 1.4 objective lens (part number = MRD31901, pixel size: 0.093 um). Cells were imaged at 100 Hz (10 ms per frame) for a total of 4 sec. To image uNS particles in *Saccharomyces cerevisiae*, Andor Yokogawa CSU-X confocal spinning disc on a Nikon Ti2 X1 microscope was used at 488 nm excitation with 10% power. Fluorescence was recorded with a sCMOS Prime 95B camera (Photometrics) with a 60x objective (pixel size: 0.18 μm), at a 100 ms image capture rate, with a time step for 1 min. To image 40nm-GEMs in U2OS mammalian cell lines, Andor Yokogawa CSU-X confocal spinning disc on a Nikon Ti2 X1 microscope was used at 488 nm excitation with 100% power. Fluorescence was recorded with a sCMOS Prime 95B camera (Photometrics) with a 60x objective (pixel size: 0.18 μm), at a 10 ms image capture rate for a total of 2 sec.

The tracking of particles was performed with the Mosaic suite of FIJI ((Sbalzarini and Koumoutsakos, 2005)), using the following parameters. For yeast 40nm-GEMs: radius = 3, cutoff = 0, Per/Abs: variable, a link range of 1, and a maximum displacement of 7 px, assuming Brownian dynamics. For yeast μNS particles: radius = 2, cutoff = 0, Per/Abs: variable, a link range of 1, and a maximum displacement of 5 px, assuming Brownian dynamics. For mammalian cell 40nm-GEMs: radius = 2, cutoff = 0, Per/Abs: variable, a link range of 1, and a maximum displacement of 5 px, assuming Brownian dynamics.

All trajectories were then analyzed with the GEM-Spa (GEM single particle analysis) software package that we are developing in house: https://github.com/liamholtlab/GEMspa/releases/tag/v0.11-beta Mean-square displacement (MSD) was calculated for every 2D trajectory, and trajectories continuously followed for more than 10 time points were used to fit with linear time dependence based on the first 10 time intervals to quantify time-averaged MSD: MSD(T)t = 4D_eff_T, where T is the imaging time interval and D_eff_ is the effective diffusion coefficient with the unit of μm^2^/s. To determine the ensemble-time-averaged mean-square displacement (MSD), all trajectories were fitted with MSD(τ)_T−ens_=4Dτ^α^ where α indicates diffusion property, with α = 1 being Brownian motion, α < 1 suggests sub-diffusive motion and α > 1 as super-diffusive motion. We used the median value of D_eff_ among all trajectories to represent each condition, and perform normalization to time point 0 min in most of the yeast experiments. For mammalian cell quantification of D_eff_, the same field of the view of the cytosolic region from individual cell was cropped before and after drug treatment, and the median value of D_eff_ from all trajectories within the cropped region were quantified, and D_eff_ from the same cell after drug treatment was normalized to before drug treatment D_eff_ as a fold change value.

Step sizes for all trajectories were extracted from GEM-Spa, to reflect the distance (with the unit of μm) of particles traveled for a given time interval: 100 ms for 40nm-GEMs and 1-sec for μNS particles. To measure the average intensity of μNS particles from GEM-Spa, a fixed radius (radius=3) was used along the movie series, and the mean intensities of particles were measured at all the tracked frames and then summarized as average mean intensity.

### Live yeast cell ATP sensor and pH sensor imaging and quantification

To image the ratiometric ATP sensor QUEEN in *Saccharomyces cerevisiae*, TIRF Nikon TI Eclipse microscope was used with 100x Phase, Nikon, oil NA = 1.4 objective lens (part number = MRD31901) at around 25°C. Both sensors were illuminated with LED light sources at a single z-plane using DAPI filter set (value recorded as 410ex): excitation filter (Excitation wavelength/ Bandwidth (FWHM) = 401/17 nm) and an emission filter (Emission wavelength/ Bandwidth (FWHM) = 444/58 nm), GFP filter set (value recorded as 480ex): excitation filter (Excitation wavelength/ Bandwidth (FWHM) = 470/40 nm) and an emission filter (Emission wavelength/ Bandwidth (FWHM) = 525/50 nm). The quantification of QUEEN sensor ratio is mainly followed by standard protocol (Takaine, 2019). Basically, individual yeast cells were segmented with 200×200 pixels ROI and the average intensity was measured after background subtraction. And the QUEEN sensor ratio was calculated as 410ex/480ex, where a reducing ratio indicates a decrease of intracellular ATP level.

To generate a calibration curve with pH sensor pHurion integrated into the yeast cells (LH4820), we used the McIlvaine buffer to prepare extracellular medium with buffer pH ranging from 5-8, mainly by varying the amount of 0.2M disodium phosphate and 0.1M citric acid. Yeast cells were permeabilized by 100 μg/ml digitonin (Sigma-Aldrich Cat. no. D141-100MG) while incubating with various pH buffers, and 20 mM 2-DG, 10 μM antimycin A. And the pH sensor pHurion was imaged as the same as ATP sensor QUEEN, the sensor ratio was calculated as 410ex/480ex for each measured pH. A linear model was fitted to determine the standard curve, so that the intracellular pH can be quantified upon acute glucose starvation.

### Live yeast cell RNP granules and Q-bodies imaging and quantification

Yeast cells expressing P-bodies protein marker (Dcp2-mScarlet), stress granules protein marker (Pab1-mNeoGreen) or Q-bodies protein marker (Hsp42-mScarlet) were immobilized on the concanavalin A precoated 384 glass bottom imaging plate. And yeast cells were imaged on TIRF Nikon TI Eclipse microscope with 100x Phase, Nikon, oil NA = 1.4 objective lens (part number = MRD31901) at around 25°C, LED light were used to illuminate the preselected field of views, and fluorescence intensity were recorded using GFP and RFP filter set. Acute glucose starvation mediums with/without cycloheximide were switched before imaging. And time lapse movies were set up to record every 10 min for a total of 120 min. 6-μm Z stacks of yeast cells were taken with 0.5-μm steps between frames. The Z stacks were projected using average intensity in Fiji, trackmate were used to identify P-body condensates or Q-bodies in individual cells, and number of P-bodies or Q-bodies were tracked along the time series.

### Immunofluorescence

U2OS cells were seeded in the 24-well glass bottom plate (Cellvis, Cat. no. P24-1.5H-Nor) to reach around 60-70% confluency the next day. After 30 min treatment with the indicated protein translation inhibitors, cells were immediately fixed with flesh 4% paraformaldehyde (Electron Microscopy Sciences, Cat. No. 15714) for 10 minutes at room temperature. The cells were subsequently washed three times with 1x PBS, and permeabilized with 0.2% Triton X-100 (Fisher Scientific, Cat. No. 9002-93-1) in 1x PBS for 15 min at room temperature. And then cells were blocked with 1% BSA, in PBST (PBS+ 0.1% Tween 20) for 1-hour before applying the primary antibodies (1:250 dilution was applied for both anti-G3BP1 and anti-Dcp1a antibodies) for overnight incubation in 4°C. The next day, solutions were removed and the cells were washed four times with 1x PBS, and incubated with secondary antibodies (1:1000 dilution was applied for both Anti-Rabbit IgG (H+L) and Anti-Mouse IgG (H+L)) in 1% BSA, in PBST at room temperature for 1-hour in the dark. Afterwards, solutions were removed and cells were washed with 1x PBS for four times before staining with 1 μM Hoechst 33342 (Thermo Fisher Scientific 62249) for 15 min at room temperature in the dark and were subsequently stored in 1x PBS at 4°C in the dark until imaging. The fixed plates of cells were imaged on an Andor Yokogawa CSU-X confocal spinning disc on a Nikon TI Eclipse microscope at room temperature. The fluorescence signals were obtained using DAPI epifluorescence (excitation wavelength/bandwidth: 395/25 and emission wavelength/bandwidth: 460/50), RFP laser (Coherent, filter: ET605/70m) and far red (Coherent, ET700/75m) lasers, and images were captured using a Prime 95B scMOS camera (Photometrics) with a 60x/1.49 numerical aperture objective lens. 8-μm Z stacks of G3BP1 and Dcp1 fluorescence were taken with 0.5-μm steps between frames. The Z stacks were projected using maximum intensity in Fiji and P-bodies numbers (Dcp1 foci) were manually counted in each cell.

### Poly-dT FISH Imaging

U2OS cells were seeded in the 24-well glass bottom plate (Cellvis, Cat. no. P24-1.5H-Nor) to reach around 60-70% confluency the next day. After 30 min treatment with 10 nM DMDA PatA, cells were immediately fixed with flesh 4% paraformaldehyde for 10 minutes at room temperature. The cells were subsequently washed three times with 1x PBS, and permeabilized with 0.2% Triton X-100 in 1x PBS in room temperature for 15 min. And then cells were washed two times with 1x PBS and one time with 2x SSC (300 mM Sodium chloride, 30 mM Sodium citrate pH8). Pre-hybridize the cells with the 200 μl pre-warmed (37°C) hybridization buffer (Molecular instruments, HCR hybridization buffer) in a 37°C humidified chamber for 30 min. 25 ng of poly-dT-alexa647 probe was prepared for each sample in the HCR hybridization buffer, and applied it to each well. And the plate was kept in a 37°C humidified chamber in the dark overnight. The next day, the hybridization buffer was removed and cells were washed twice vigorously with a pre-warmed HCR washing buffer (Molecular instruments), followed by an additional 2 times wash of 1x PBS. 1 μM Hoechst 33342 was applied into each well and incubated in dark for 15 min in 1x PBS to stain the nucleus. Cells were subsequently stored in 1x PBS at 4°C in the dark until imaging. The fixed plates of cells were imaged on an Andor Yokogawa CSU-X confocal spinning disc on a Nikon TI2 X1 microscope at room temperature. The fluorescence signals were obtained with DAPI (Coherent, filter: ET455/50m) and far red (Coherent, ET700/75m) lasers, and images were captured using a Prime 95B scMOS camera (Photometrics) with a 60x/1.49 numerical aperture objective lens. 10-μm Z stacks of poly-dT FISH fluorescence were taken with 0.2-μm steps between frames.

### Blue-light activation of optogenetic artificial RNP condensates

Two days post transient transfection of pLH1972 or pLH2126 into U2OS 40nm-GEMs cells in the 6-well glass bottom plate, both CRY2-mCherry and CRY2-PUM.HD-mCherry proteins were expressed as diffusive pattern in the cytoplasm. On the day of the experiment, cells were mounted on a Nikon TI2 X1 spinning disk confocal scanning microscope, equipped with a 60x/1.49 numerical aperture objective lens and incubator to maintain 37°C and 5% CO2. Individual cells with CRY2-mCherry/CRY2-PUM.HD-mCherry expression were pre-selected initially, and 40nm-GEMs movies were recorded as previously described in the method section, which reflects macromolecule diffusivity before light induced artificial pumilio granule formation. To apply blue light to induce artificial pumilio granule formation, GFP laser (Coherent, filter: ET525/36m) with 25% power were illuminated to the field of view every 5 sec for 5 min, and artificial RNP condensates signal were recorded by RFP laser (Coherent, filter: ET605/70m). Afterwards, 40nm-GEMs movies were recorded again.

Graphic illustrations were created with BioRender.com.

## Acknowledgements

We thank Sarah Keegan for building the GemSpa Python package for single particle analysis. We thank Stephanie Patchett for help with the polysome profiling experiments. We thank Zoher Gueroui, Emily M. Sontag for helpful discussion and plasmids sharing. We thank Masakatsu Takaine for sharing ATP sensor plasmid. We thank J. Paul Taylor for sharing U2OS mammalian cell lines. We thank Daniel Romo for sharing des-methyl, des-amino pateamine A. We thank Tong Shu, Marie-Christin Spindler, Julia Mahamid, Alexandros Papagiannakis for helpful discussions. We thank all members in the Holt lab for their support and discussions. We thank all members in the Gresham lab for helpful discussion and manuscript edits. Figures 2B, 3A, 4C, 5A, 6 and supplementary figure 1B were created partly using Biorender. This work was supported by the NIGMS award R01GM107466 (DG), and L.J.H. was funded by NIH R01 GM132447, R37 CA240765, the American Cancer Society Cornelia T Bailey Research Award, the NIH Director’s Transformative Research Award TR01 NS127186, the Air Force Office of Scientific Research (AFoSR FA9550-21-1-3503 0091), and the Human Frontier Science Program (RGP0016/2022-102.)

---

**Supplementary figure 1.**
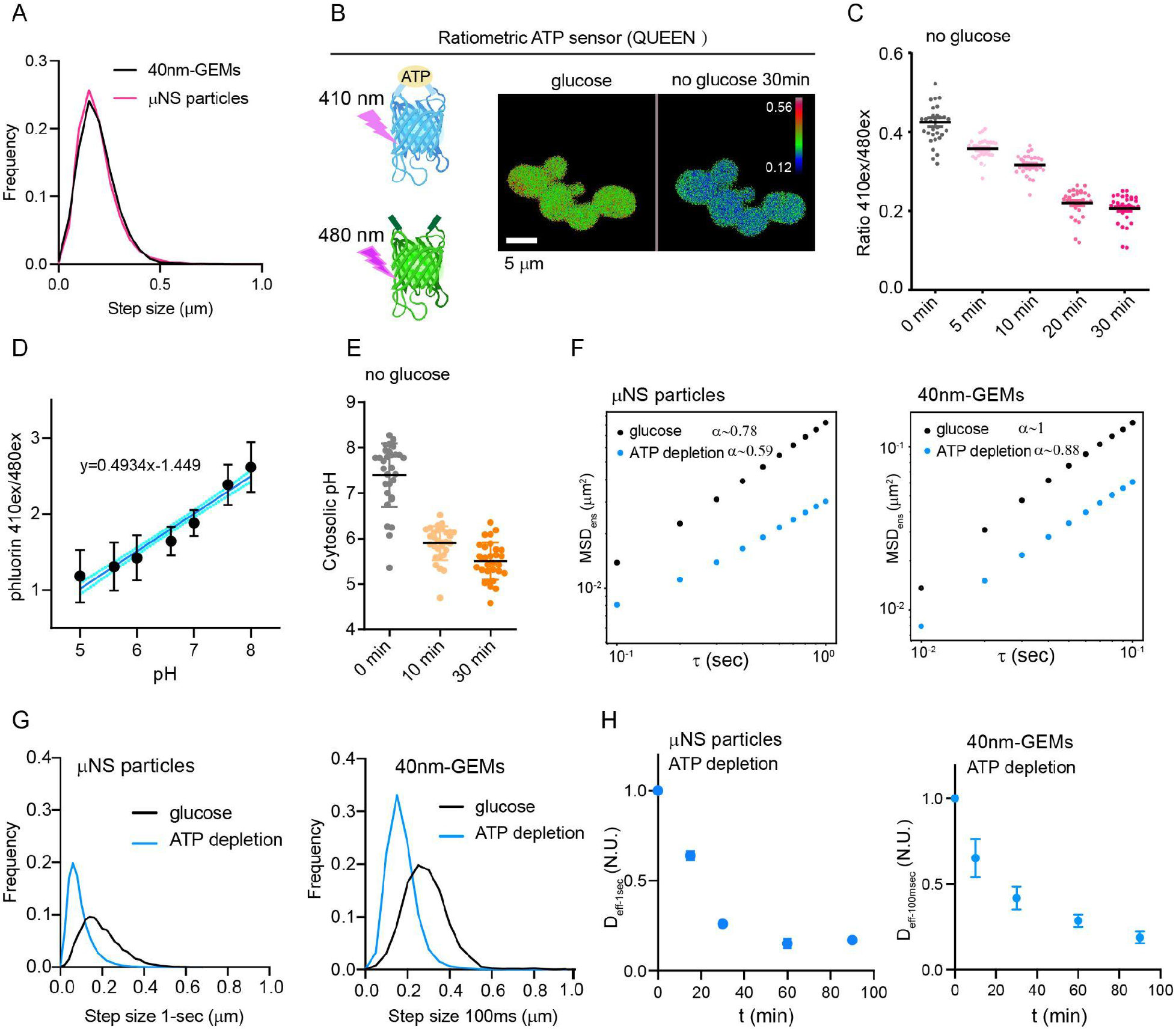
ATP reduction and cytoplasmic acidification decrease mesoscale macromolecular diffusivity, related to Figure 1. **A**. Step size distribution of 40nm-GEMs in 100 ms and μNS particles in 1 second. **B.** Graphic illustration of QUEEN sensor. ATP QUEEN sensor displays a bimodal excitation spectrum with peaks at 410 nm (ATP-bound) and 480 nm (ATP-unbound) and an emission maximum at 510 nm. Representative QUEEN ratio (410 nm ex/480 nm ex) images of yeast cells in glucose rich and glucose depleted conditions. The QUEEN ratio is pseudo-colored. **C.** Time course analysis of the QUEEN ratio after glucose depletion (n=30 cells, mean±SEM). **D.** pH calibration curve determined using permeabilized *S. cerevisiae* cells expressing cytosolic pHluorin sensor. A linear model was fitted to determine the standard curve (n=30 cells for each pH condition). **E.** The intracellular pH was quantified based on the standard curve of pHluorin sensor in *S. cerevisiae* cells upon acute glucose starvation (n=30 cells, mean±SEM). **F.** Ensemble-averaged mean-squared displacement (MSD) versus time delay (τ), log10 scale. A linear model was fitted to determine the anomalous exponent α values for glucose rich (glucose) or ATP depleted (ATP depletion) for μNS particles and 40nm-GEMs at 30 minutes (ATP depleted condition: synthetic complete medium buffer at pH 5.5 without glucose, 20 mM 2-deoxyglucose and 10 μM antimycin A and 80 mM sorbitol were supplemented. Trajectories analyzed for μNS particles, glucose: n=16373 trajectories; ATP depletion: n=16373 trajectories; For 40nm-GEMs, glucose: n=2620 trajectories; ATP depletion: n=3176 trajectories). **G.** Step size distribution of μNS particles in one second time scale, 40nm-GEMs in 100 msec time scale in glucose rich and ATP depleted conditions at 30 minutes (For μNS particles, glucose: n=16373 trajectories; ATP depletion: n=16373 trajectories; For 40nm-GEMs, glucose: n=2620 trajectories; ATP depletion: n=3176 trajectories). **H.** Fold change of median effective diffusion coefficients at one sec time scale for μNS particles, and 100 msec (D_eff-100ms_) for 40nm-GEMs from 3 biological replicate experiments (mean ± SEM.) upon ATP depletion conditions in *S. cerevisiae* cells.

**Supplementary figure 2.**
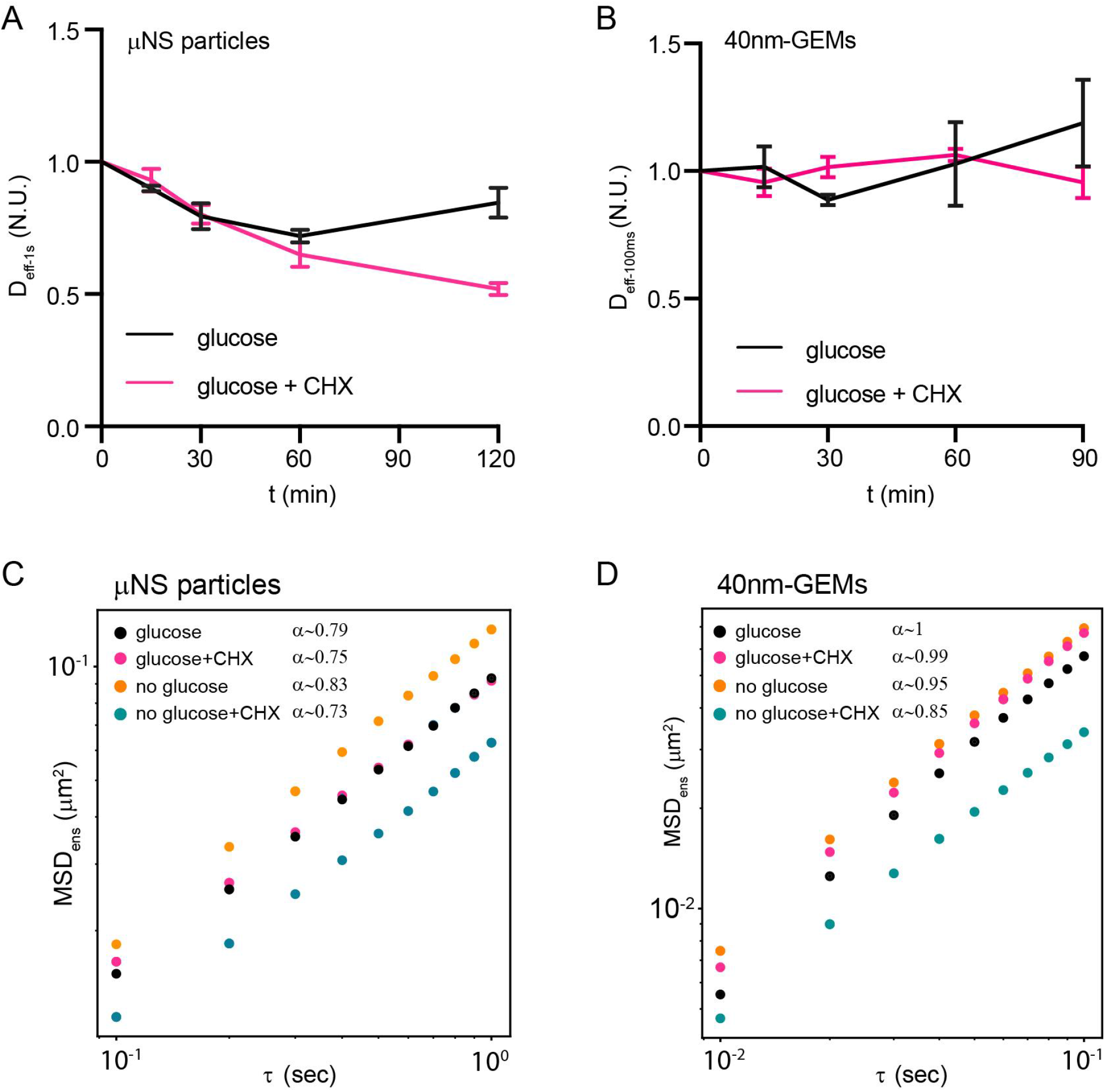
Mesoscale μNS particles and 40nm-GEMs dynamics in wild type *S. cerevisiae* cells, related to figure 2. **A.** Fold change of median effective diffusion coefficients for μNS particles from 3 biological replicate experiments in the indicated condition (CHX: cycloheximide, mean ± SEM.). **B.** Fold change of median effective diffusion coefficients for 40nm-GEMs from 3 biological replicate experiments in the indicated conditions (CHX: cycloheximide, mean ± SEM.). **C.** Ensemble-averaged mean-squared displacement (MSD) versus time delay (τ), log10 scale. A linear model was fit to determine the anomalous exponent α values for μNS particles in the indicated conditions at 30 minutes (n>10000 trajectories in each condition, CHX: cycloheximide). **D.** Ensemble-averaged mean-squared displacement (MSD) versus time delay (τ), log10 scale. A linear model was fit to determine the anomalous exponent α values for 40nm-GEMs in the indicated condition at 30 minutes (n>5000 trajectories in each condition, CHX: cycloheximide).

**Supplementary figure 3.**
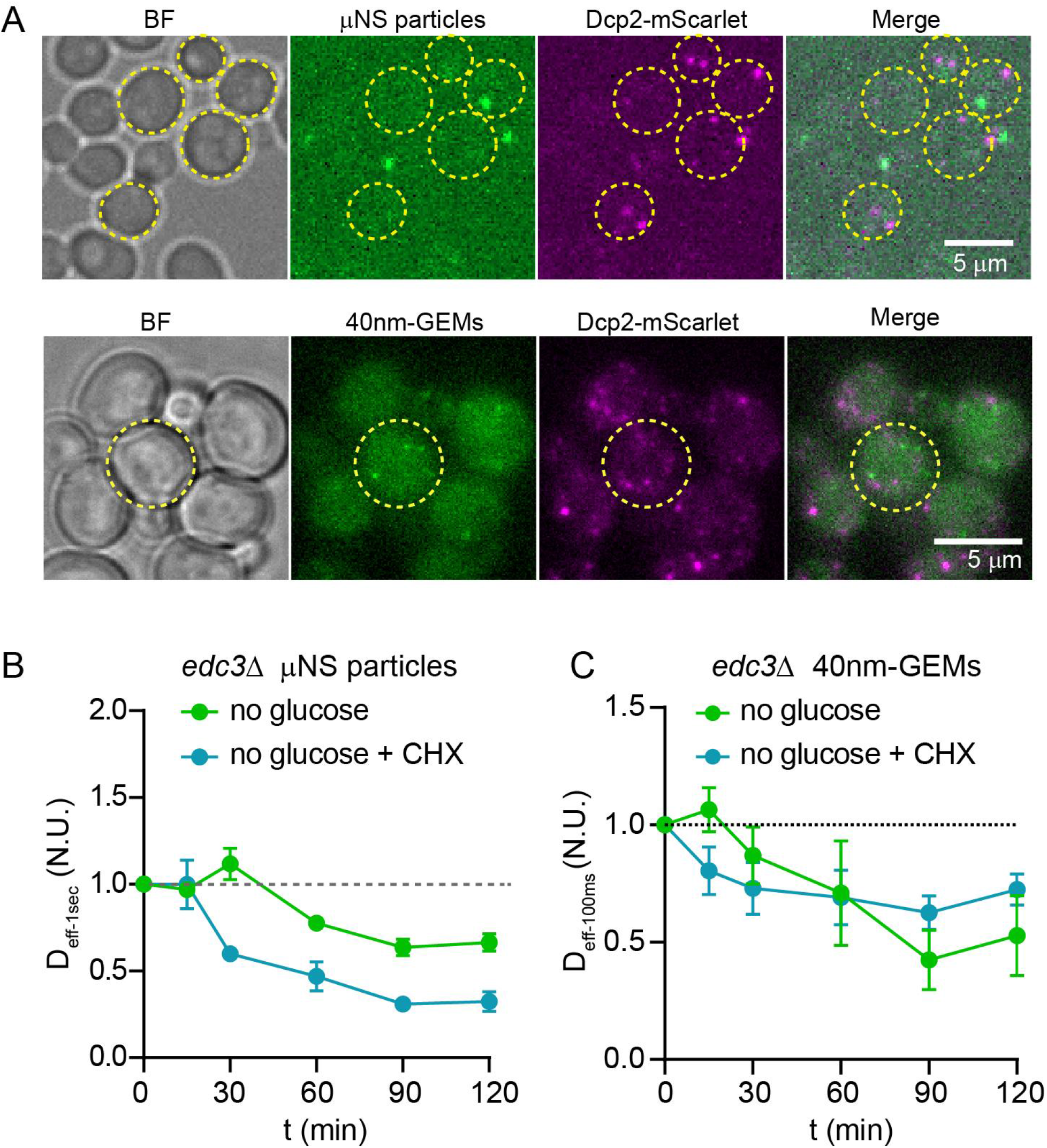
Mesoscale μNS particles and 40nm-GEMs behavior in *edc3Δ S. cerevisiae* cells, related to figure 3. **A.** P-bodies (Dcp2-mScarlet marker) are not colocalized with μNS particles or 40nm-GEMs (BF = Bright field). **B.** Fold change of median effective diffusion coefficients for μNS particles from 3 biological replicate experiments at the indicated conditions (CHX: cycloheximide, mean ± SEM.). **C.** Fold change of median effective diffusion coefficients for 40nm-GEMs from 3 biological replicate experiments at the indicated conditions (CHX: cycloheximide, mean ± SEM.).

**Supplementary figure 4.**
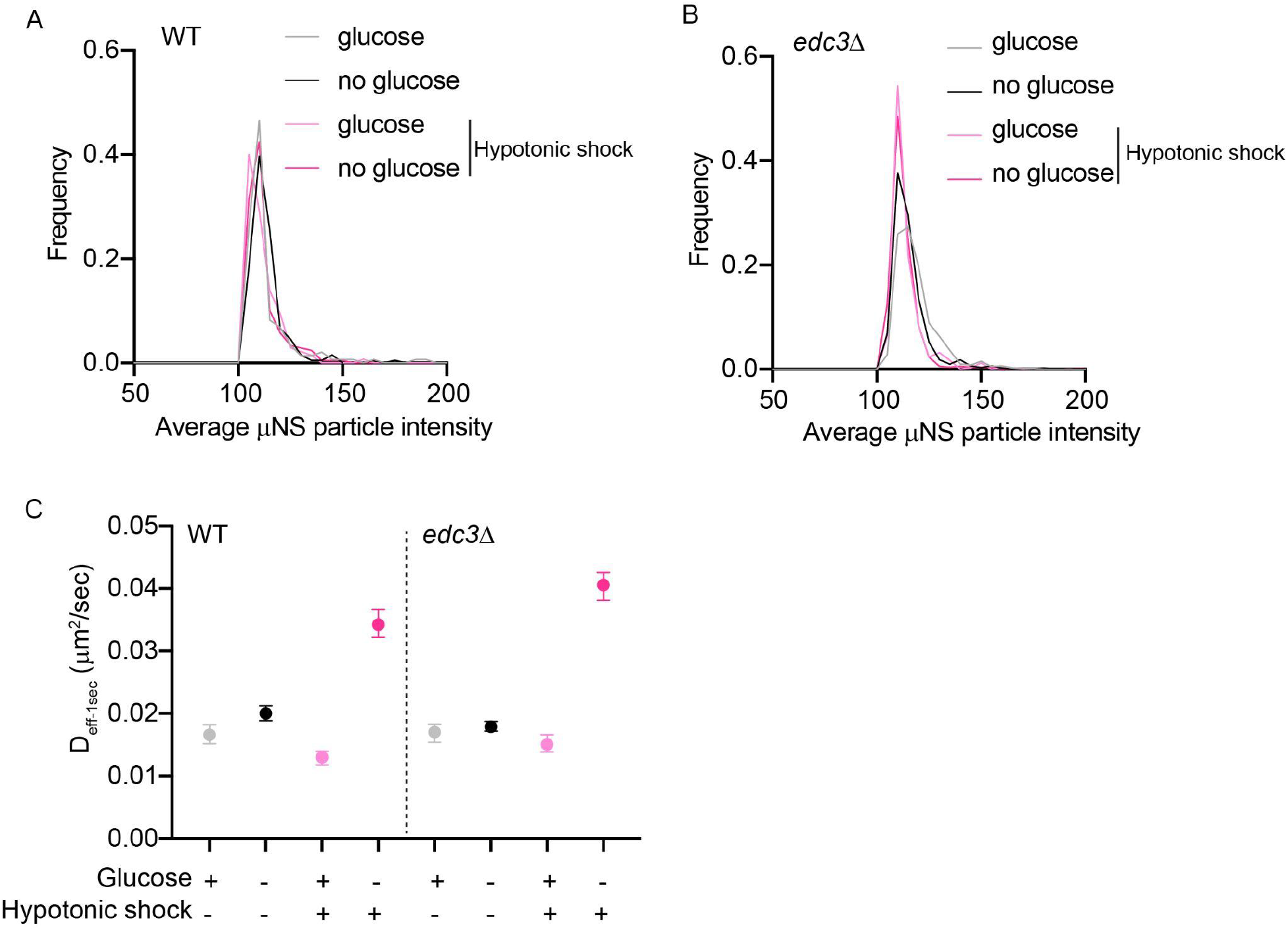
Mesoscale μNS particles diffusivity increases upon hypotonic shock, related to figure 4. **A.** Distribution of average particle intensities for µNS particles in the indicated conditions (n>700 trajectories in each condition) for wildtype and **B.** *edc3Δ* cells. **C.** Effective diffusion coefficients at 1-second time scale for μNS particles in the indicated conditions in wildtype and *edc3Δ* cells. All glucose starvation results were assessed 5 min after starvation (n>700 trajectories in each condition, median ± 95% confidence interval).

**Supplementary figure 5.**
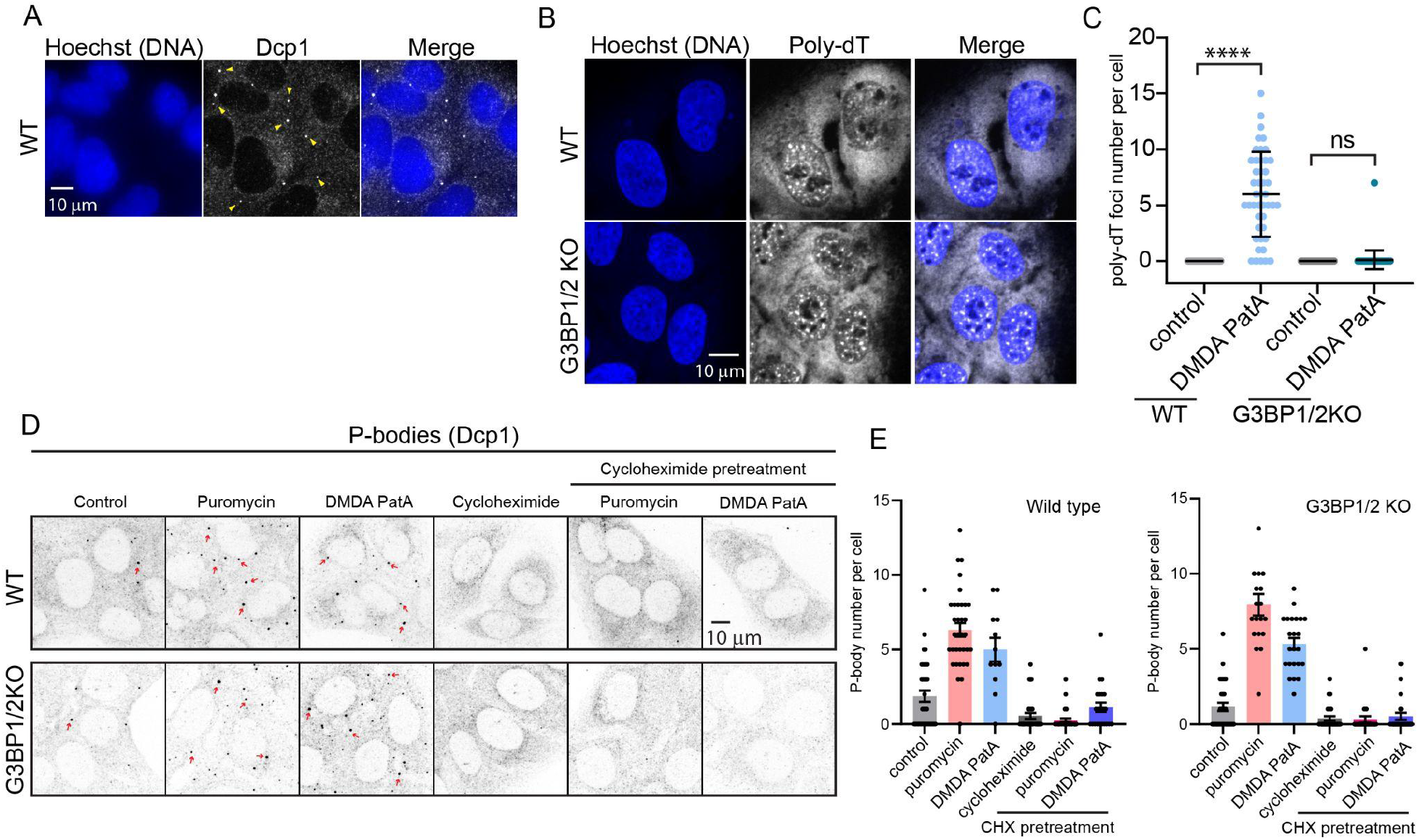
Characterization of P-bodies and mRNA localization upon treatment with translation inhibitors, related to figure 5. **A.** Representative immunofluorescence images of P-bodies (Dcp1) in U2OS wild type cells in the absence of stress. Hoechst dye was used to stain DNA. **B.** Representative poly-dT FISH images for poly(A) RNAs in wild type and G3BP1/2 KO U2OS cells without any drug treatment. **C.** Quantification of poly-dT FISH foci number in the cytoplasm with the indicated conditions (WT control: n=57 cells; WT DMDA-PatA n=45 cells; G3BP1/2 KO control n=76 cells; G3BP1/2 KO DMDA-PatA n=67 cells, mean ± SD. Statistical comparison was by a one-way ANOVA test. ns=not significant, *p<0.05, **p<0.01, ****p<0.0001). **D.** Representative immunofluorescence images of P-bodies (Dcp1) in U2OS wild type and G3BP1/2KO cells. **E.** Quantification of Dcp1 foci (P-bodies) number in U2OS wild type and G3BP1/2KO cells with the indicated translation inhibitors (WT control n=35; WT puromycin n=34; WT DMDA-PatA n=12; WT cycloheximide n=32; WT Cycloheximide pretreatment puromycin n=35; WT Cycloheximide pretreatment DMDA-PatA n=22. G3BP1/2 KO control n=31; G3BP1/2 KO puromycin n=19; G3BP1/2 KO DMDA-PatA n=23; G3BP1/2 KO cycloheximide n=28; G3BP1/2 KO Cycloheximide pretreatment puromycin n=24; G3BP1/2 KO Cycloheximide pretreatment DMDA-PatA n=28. mean ± SD. Statistical comparison was by a one-way ANOVA test. ns=not significant, *p<0.05, **p<0.01, ****p<0.0001).

